# Pan-genome and multi-omics analyses reveal the genomic basis of symbiotic evolution in the mycoheterotrophic medicinal orchid *Gastrodia elata*

**DOI:** 10.64898/2026.01.26.701689

**Authors:** Mingjin Huang, Shanshan Luo, Linshuang Tang, Hao Yin, Daliang Liu, Zhipeng Li, Qiyu Chen, Yinjie Jiao, Mengge Li, Yanlin Hao, Tao Li, Dachang Wang, Hongchang Liu, Dandan Li, Jin He, Lin Cheng, Cheng Li, Hualei Wang, Guojin Zhang, Wei Wang, Ruimin Wang, Xinhao Sun, Yiyong Zhao

## Abstract

Mycoheterotrophy represents an extreme evolutionary strategy in which plants abandon photosynthesis and become obligately dependent on fungal partners for carbon and nutrients. *Gastrodia elata*, a medicinal orchid forming long-term symbioses with *Armillaria* and *Mycena*, provides a compelling system to investigate the genomic basis of obligate plant-fungus mutualism. Here, we generate a chromosome-level genome assembly (1.09 Gb) of a dark-red *G. elata* accession and construct a pan-genome from 12 *Gastrodia* accessions to dissect the evolutionary consequences of mycoheterotrophy. Phylogenomic analyses reveal pervasive hybridization across cultivated germplasm and demonstrate that tuber morphology more accurately reflects intraspecific relationships than stem color, challenging the traditional color-based taxonomy. Comparative genomic analyses reveal a pronounced degeneration of photosynthetic capacity. We detect no intact nuclear-encoded *rbcS* loci, which would preclude assembly of the Rubisco holoenzyme and thus canonical Calvin-Benson carbon fixation. Notably, a subset of photosynthesis-related genes is retained and transcriptionally active, suggesting their co-option into non-photosynthetic regulatory roles. Despite millions of years of intimate association with fungal partners, we detect no evidence of fungal-derived horizontal gene transfer based on a multi-step phylogenomic validation pipeline, indicating that *G. elata*-fungus mutualism is sustained through metabolic exchange rather than genetic integration. We further identify the *Gastrodia* antifungal protein (GAFP) gene family as a key molecular innovation underlying symbiotic homeostasis. The recently derived Class 1 GAFP lineage exhibits distinct domain architecture, tandem expansion, and elevated expression, consistent with enhanced regulation of fungal overgrowth. Finally, we delineate a specialized nutrient-acquisition framework adapted to obligate heterotrophy, in which fungal trehalose hydrolysis supplies hexose carbon via an expanded sugar transporter protein (STP) repertoire, while organic nitrogen is assimilated through amino acid and oligopeptide transporters coupled with urease-mediated conversion, compensating for the loss of nitrate uptake and reduction pathways. Together, these results establish a pan-genomic blueprint for symbiotic adaptation and reductive genome evolution in *G. elata*.

## Introduction

*Gastrodia elata* Blume (GEB), commonly known as Tian-ma in Chinese, is a mycoheterotrophic perennial herb thriving in the humid mountainous regions of East Asia^1, 2^. The *Flora of China* recognizes several intraspecific forms largely defined by stem and floral coloration, including red, green, brown and yellow phenotypes, as well as the locally named ‘Song Tian-ma’ form (Figure 1A), with the red variant designated as the original form (*G. elata* f. *elata*). Red *G. elata*, brown *G. elata* and their hybrids, are especially valuable as traditional Chinese medicinal materials. After ∼50 years of cultivation and breeding, including frequent inter-form crosses in China, *G. elata* has developed significant diversity in germplasm resources. However, previous studies on *G. elata* have focused on color, limiting the understanding of its reproductive strategies and the incomplete analysis of its genetic diversity^3–6^. Existing genomic studies have predominantly focused on brown and green variants, while genomic information for red *G. elata* remains notably scarce despite its widespread cultivation as a major commercial variety. This gap limits a comprehensive view of intraspecific diversity. It also leaves the genomic status of the red form—often regarded as the ‘original’ form—largely untested.

**Figure 1.**
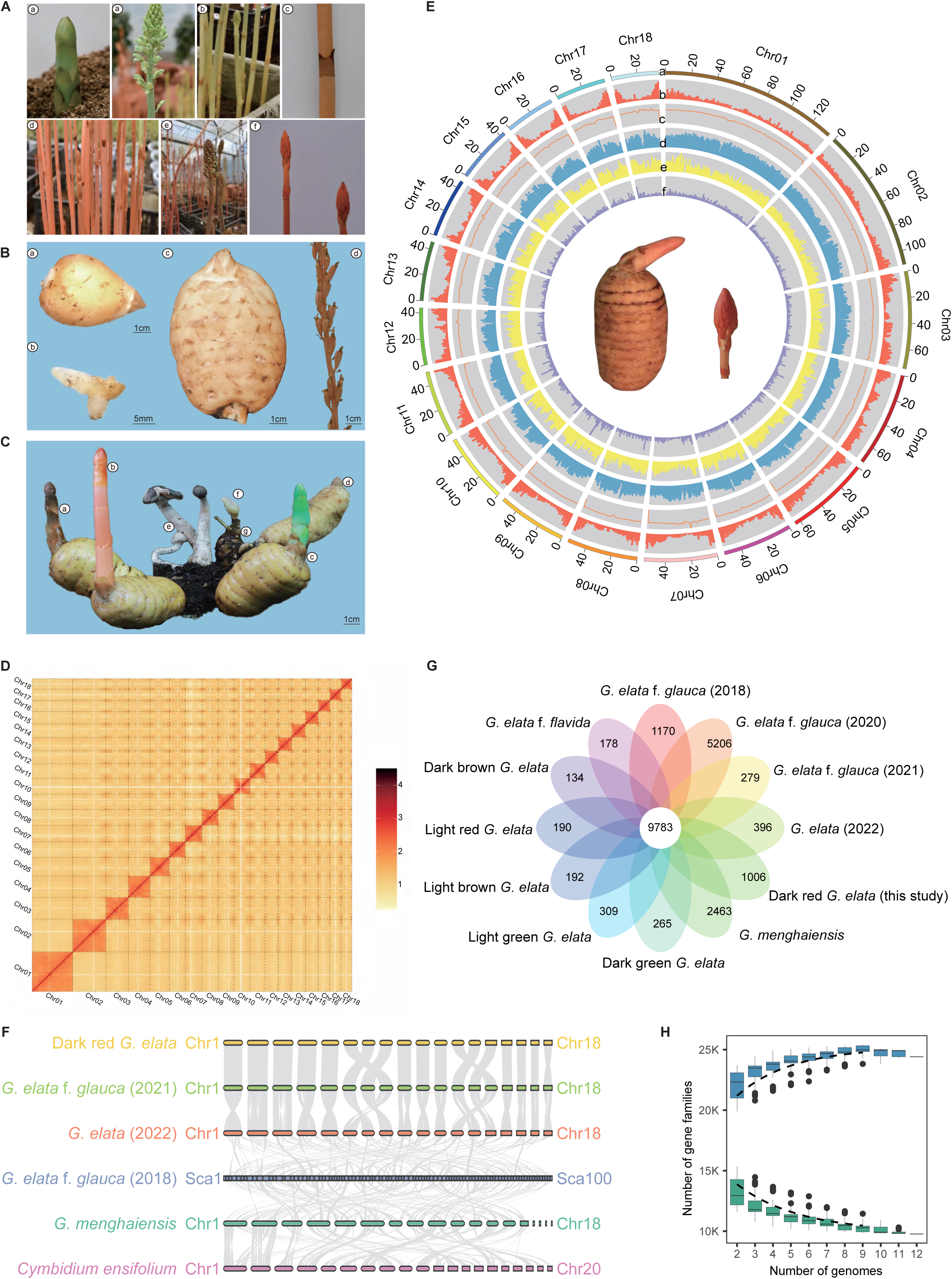
Morphological features, genome sequencing assembly, and pan-gene analysis of *G. elata*. A. Morphological traits of stems, inflorescences, and tubers in seven cultivated *G. elata* genotypes. a. dark green *G. elata*, b. light green *G. elata*, c. dark-red G*. elata*, d. light red *G. elata*, e. dark brown *G. elata,* f. light brown *G. elata*, g. *G. elata* f*. flavida*. B. Five developmental stages of dark-red G*. elata* f. *elata*: a. immature tuber (TMMM), b. developing tuber (TMBM), c. mature tuber (TMJM), d. stem and flower (TMHJ), and e. capsules (TMGS). C. Schematic diagram of the symbiotic relationship between *G. elata* and *Armillaria gallica*. a-d. four morphotypes of *G. elata* with dark brown, dark red, dark green, and light red coloration respectively; e. *Armillaria gallica*; f. newly formed tuber; g: symbiotic tuber. D. Hi-C interaction heatmap of 18 chromosomes in the dark-red G*. elata* genome E. A Circos plot illustrating the genomic features of dark-red G*. elata* distributed across 18 chromosomes. a. chromosomes; b. gene density; c. GC content; d-f. Gypsy, Copia, and DNA transposable element densities. F. Synteny plot of *Cymbidium ensifolium*, *G. menghaiensis*, and four *G. elata* genomes, with gray lines indicating syntenic blocks. Chr: chromosome; Sca: scaffold. G. Venn diagram comparing the genomes of *G. menghaiensis* and 11 *G. elata* species, displaying the number of shared and unique gene families. H. Counts of pan-gene (blue) and core-gene (green) with increased samples.

As a fully heterotrophic plant that has completely lost photosynthetic capability, *G. elata* exhibits a unique survival strategy. Due to the lack of endosperm in its seeds, *G. elata* must first establish a symbiotic relationship with fungi of the genus *Mycena* for germination, and then transition to symbiosis with *Armillaria mellea* for subsequent development (Figure 1B, C), which provides carbon and nutrition sources essential for its energy metabolism^7^. This long-term mycoheterotrophic lifestyle has driven the loss of genes associated with photosynthesis and root development, while simultaneously inducing significant specialization in floral architecture—the sepals fuse to form a cylindrical perianth tube, and anthers remain enclosed by persistent caps and do not dehisce autonomously, necessitating artificial pollination^8, 9^.

Notably, the symbiotic relationship between *G. elata* and *Armillaria* represents a complex evolutionary trade-off. On one hand, *G. elata* depends on *Armillaria* for nutrient acquisition; on the other hand, excessive infection by *Armillaria* can lead to tuber rot and severely reduced yield^10, 11^. To cope with this challenge, *G. elata* has evolved a unique defense mechanism—the antifungal protein (GAFP) extracted from tubers can inhibit excessive growth of *Armillaria* fungi, thereby establishing a defensive system^12^. However, the composition of the gene family GAFP, its molecular mechanisms of action, interactions with Armillaria proteins, and evolutionary trajectory during adaptation to mycoheterotrophy remain largely unknown.

This intimate symbiotic relationship spanning millions of years naturally raises another important scientific question: has horizontal gene transfer (HGT) occurred between *G. elata* and its fungal partners? Recent studies have demonstrated that plant-microbe symbiotic systems may represent “hot spots” for HGT^3, 13–16^. The *G. elata*–fungus *(Armillaria* spp./*Mycena* spp.) symbiotic system provides an ideal model for studying interkingdom HGT^5, 17, 18^. However, whether symbiosis drives functional gene transfer and how exogenous genes integrate into host genomes remain unresolved. At the level of nutrient transport mechanisms, genes related to sucrose transporters (SUTs) and ammonium transporters (AMTs) have been identified in *G. elata*, providing molecular evidence for absorption and utilization of carbon and nitrogen nutrients from symbiotic fungi. Ho et al. identified a functional sucrose-specific transporter *GeSUT4*, suggesting sucrose may be the main sugar transported between *G. elata* and *Armillaria*^19, 20^. However, critical questions remain: Is the sugar obtained from *Armillaria* limited to sucrose, or does it include other types? What are the transport mechanisms for amino acids and fatty acids^21, 22^? Additionally, Rubisco serves as an important photosynthetic enzyme, and the CAM pathway regulated by the *PEPC* gene family represents a carbon fixation strategy^23, 24^; however, their presence and function in *G. elata* remain unknown.

To fill the knowledge gap mentioned above, we performed whole-genome sequencing and assembly of *Gastrodia elata* f. *elata* (dark-red G*. elata*), and integrated analyses of pangenomics, comparative genomics and transcriptomics. The main research objectives are as follows: (1) Integrate published genome data of *G. elata* with newly generated sequencing data from this study to construct a pangenomic framework, and provide novel insights into the taxonomic status of *G. elata* variants; (2) construct a phylogenetic tree of species based on 54 representative green plants, and reveal the genomic evolutionary characteristics of *G. elata* through comparative genomic analysis; (3) elucidate the genetic basis underlying the tubular floral morphology of *G. elata*, and explore the adaptive evolutionary mechanisms of *G. elata* antifungal protein (GAFP) genes in defending against *Armillaria* infection; (4) investigate whether horizontal gene transfer has occurred between *G. elata* and its symbiotic fungi, and predict the types of nutrients acquired by *G. elata* from *Armillaria* and their corresponding molecular transport mechanisms. This study will lay a theoretical foundation for understanding the evolutionary adaptive mechanisms of *G. elata* as well as the conservation and utilization of its germplasm resources.

## Results

### Genome assembly, annotation and pan-genome construction

The genome of dark-red G*. elata* was sequenced using 42 Gb PacBio long reads (38.52× coverage) and 140 Gb Illumina paired end reads (Supplementary Fig. 1A and Supplementary Table 1). The assembled genome spans 1.09 Gb with a contig N50 of 53.67 Mb, surpassing all previously published *G. elata* genomes in assembly continuity and completeness (Supplementary Table 2). Hi-C scaffolding anchored 1,070,965,212 bp (98.01%) onto 18 chromosomal-level scaffolds, with 461 scaffolds remaining unplaced (Fig.1D, 1E and Supplementary Table 3). The genome has a GC content of 34.46% (Supplementary Fig.1B and Supplementary Table 4). BUSCO analysis identified 77.80% complete BUSCOs (Supplementary Fig.1C and Supplementary Table 5), and CEGMA assessment showed 84.68% complete and 93.95% complete plus partial matches to 248 core eukaryotic genes (Supplementary Table 6). Illumina reads exhibited a 99.84% mapping rate with 98.89% genome coverage (Supplementary Fig.1D and Supplementary Table 7). The heterozygous and homozygous SNP rates were 0.000572% and 0.000026%, respectively (Supplementary Table 8).

Repetitive sequences constitute 69.38% of the genome, with transposable elements accounting for 67.85% (Supplementary Table 9). When TE divergence levels are below 60%, proportions follow the order LTR > LINE > DNA (Supplementary Fig.2A), indicating recent replication and insertion events. A total of 19,477 protein-coding genes were predicted, with 90.57% (17,640) functionally annotated (Supplementary Fig.2B and Supplementary Table 10).

**Figure 2.**
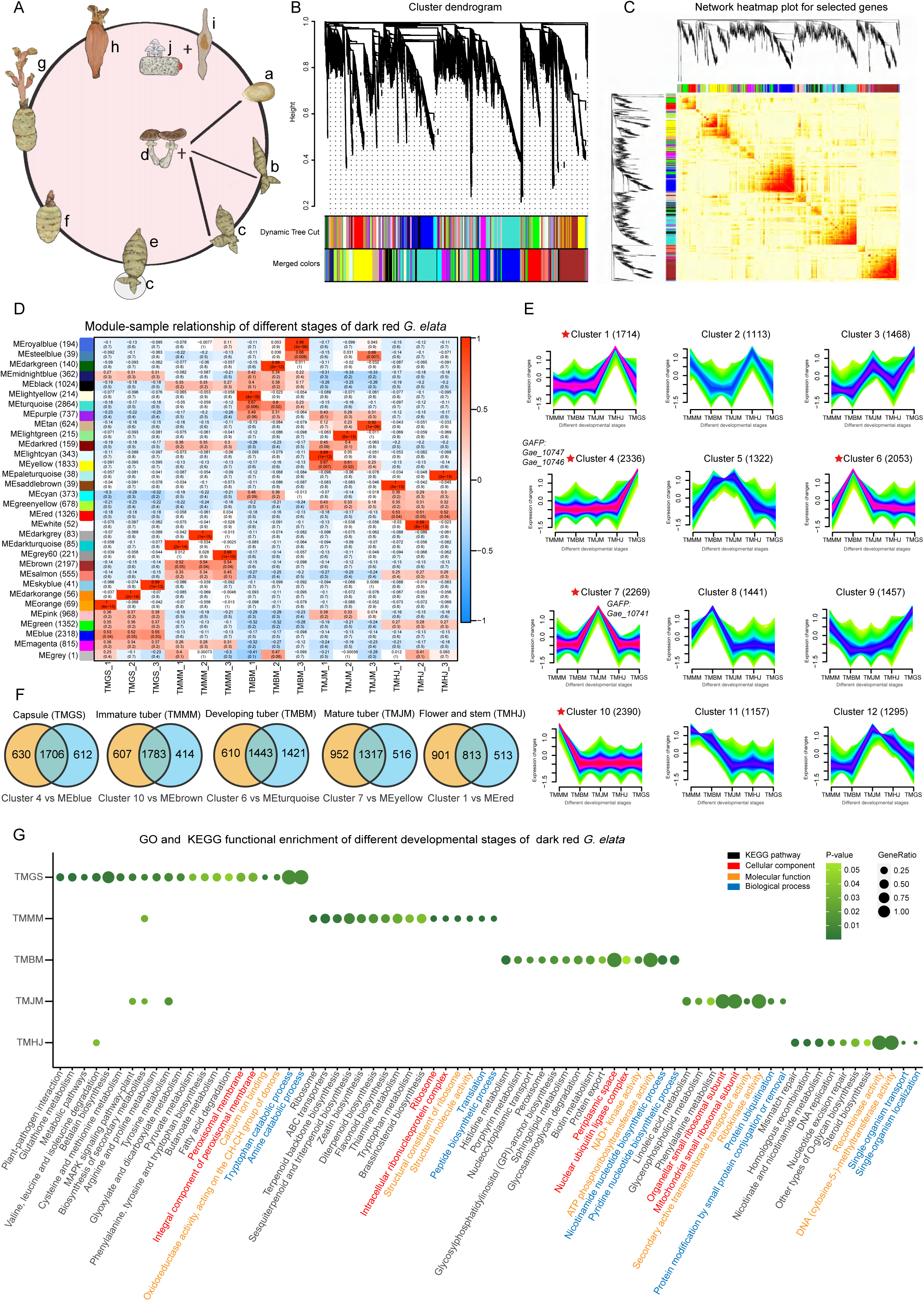
Analysis of transcriptome data for dark-red G*. elata* at various developmental stages. A. *G. elata* life cycle: a. protocorm, b. immature tuber (TMMM), c. developing/maturing tuber (TMBM), d. *Armillaria*, e. mature tuber (TMJM), f. mature tuber for reproductive growth, g. flowering mature tuber (TMHJ), h. capsule (TMGS), i. seed, j. *Mycena*. B. Weighted gene co-expression network analysis (WGCNA) dendrogram showing 31 co-expression modules. Each of the 20,757 genes corresponds to a tree tip, with each module forming a major branch. The lower panel illustrates the modules in designated colors. C. WGCNA heatmap of selected genes, with color intensity (white-yellow-red) indicating pairwise gene connectivity. D. Module-sample correlation heatmap for dark-red G*. elata* developmental stages, showing Pearson correlation coefficients colored by score. The number of genes in each module is shown on the left. E. Key co-expression module clusters across five developmental stages. Stars indicate optimal clusters. The x-axis represents developmental stages; the y-axis represents expression fold changes. The number of genes in each cluster is shown in parentheses. F. Venn diagrams showing the overlap between WGCNA module genes (D) and expression pattern clusters (E). Yellow circles represent genes in each cluster; blue circles represent genes in the corresponding module; overlapping regions indicate shared genes.G. GO and KEGG functional enrichment analysis of overlapping genes (P < 0.05). The x-axis represents developmental stages; the y-axis represents GO terms/KEGG pathways.

To explore genomic diversity, we constructed a pan-genome from 12 *Gastrodia* genomes (Supplementary Table 5). After standardized reassembly, contig N50 values ranged from 1,284 kb to 1,575 kb, BUSCO scores from 61.90% to 77.80% (dark-red G*. elata* highest). Collinearity analysis showed dark-red G*. elata* exhibited the best syntenic relationship with *G. elata* f. *glauca* (2021) (Fig.1F). Approximately 9,783 gene families constituted the core genome across all 12 samples (Fig.1G). Pan-genome size stabilized with increasing sample number, while core genome size decreased initially then plateaued (Fig.1H). GO analysis revealed non-core genes enriched in substance transport and metabolism (Supplementary Table 11-13). Despite lacking photosynthetic capacity, *G. elata* retains genes involved in photosensitivity, light response, and photosynthesis (Supplementary Table 14).

### Transcriptome profiling of *G. elata*

Gene expression distribution, Pearson correlation coefficients, and principal component analysis validated the reliability of transcriptome sequencing across stems of different colors in *G. elata* forms (Supplementary Fig. 3A, 3B, 3C). A total of 786.48 million high-quality reads were generated (Supplementary Table 15), with 12,237 shared expressed genes identified (Supplementary Fig. 3D). Differentially expressed genes (DEGs) were identified as follows: TMQG vs TMSR (9,218), TMQR vs TMSR (8,066), TMSG vs TMSR (5,292), TMQB vs TMSR (5,083), and TMSB vs TMSR (3,932) (Supplementary Fig.4). GO and KEGG enrichment analyses revealed that DEGs were significantly enriched in gene expression regulation, biosynthetic and metabolic processes, phenylpropanoid biosynthesis, fatty acid elongation, plant MAPK signaling pathway, DNA replication, and flavonoid biosynthesis (Supplementary Fig.5A, 5B). Phenylpropanoid metabolism produces over 8,000 metabolites including lignin, flavonoids, and lignans, and DEGs involved in flavonoid biosynthesis may contribute to variations in flower and stem coloration^25^. Notably, TMQG and TMQR exhibited higher DEG numbers compared to TMSR, while TMSB showed the lowest, indicating a close genetic relationship between TMSB and TMSR. This relationship was further supported by cluster analysis of 13 volatile compounds detected by GC-MS (Supplementary Fig.6 and Supplementary Table 16). Transcriptome sequencing of five developmental stages of dark-red G*. elata*, including capsule (TMGS), immature tuber (TMMM), developing tuber (TMBM), mature tuber (TMJM), and inflorescence (TMHJ) (Fig. 2A, a, b, c, g, h), identified 20,015 highly expressed genes. Weighted Gene Co-expression Network Analysis (WGCNA) classified these genes into 32 stage-specific modules with distinct expression profiles (Fig. 2B, 2C, 2D). Stage-specific highly expressed genes numbered 2,318 (TMGS), 2,197 (TMMM), 2,864 (TMBM), 1,833 (TMJM), and 1,326 (TMHJ) (Fig. 2D). Time-series clustering further classified these genes into 12 clusters associated with specific developmental stages (Fig. 2E). Comparison of WGCNA and time-series analysis revealed substantial gene overlap in vegetative stages: 1,706 (TMGS), 1,783 (TMMM), 1,433 (TMBM), and 1,317 (TMJM) shared genes (Fig. 2F). In contrast, the flowering stage (TMHJ) displayed only 813 shared genes, suggesting more specialized gene expression dynamics during reproductive development. GO and KEGG analyses of capsule-specific genes highlighted involvement in plant-pathogen interaction, glutathione metabolism, and amino acid degradation. Two antifungal protein-coding genes were highly expressed in capsules, suggesting potential antifungal properties (Fig. 2E, Cluster 4). Highly expressed genes in other stages were enriched in substance transport and biosynthesis, potentially crucial for tuber growth and active ingredient formation (Fig. 2G).

**Figure 3.**
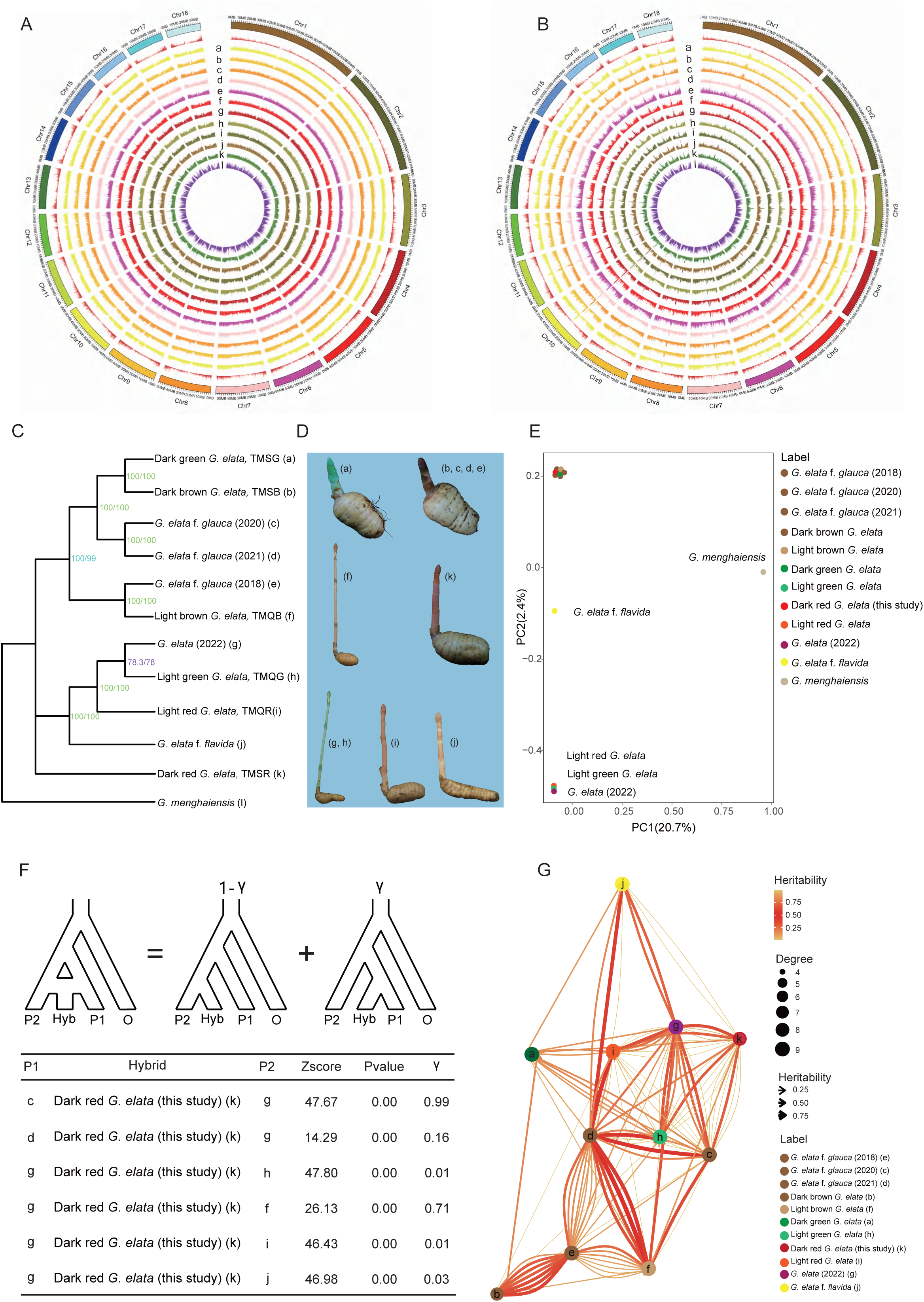
Phylogeny and hybridization analysis of *Gastrodia* species A-B. Circos plots show SNPs (A) and INDELs (B) among 12 *Gastrodia* plants, including 11 *G. elata* and one *G. menghaiensis*. Outer ring: chromosome length; second ring: dark-red G*. elata* gene density. a. Dark green *G. elata*; b. Dark brown *G. elata*; c. *G. elata* f. *glauca* (2020); d. *G. elata* f. *glauca* (2021); e. *G. elata* f. *glauca* (2018); f. Light brown *G. elata*; g. *G. elata* (2022); h. Light green *G. elata*; i. Light red *G. elata*; j. *G. elata* f. *flavida*; k. dark-red G*. elata* (this study); l. *G. menghaiensis*. C. Phylogenetic tree of 11 *G. elata* forms and one *G. menghaiensis*. D. Oval tubers (samples a-f, k) and elongated tubers (samples g-j) of *G. elata* were used in this study. E. PCA of 12 *Gastrodia* SNPs datasets visualizes sample clustering. F. Hybridization analysis of cultivated dark-red G*. elata* inferred by HyDe (Hybrid Detection) software. G. Hybrid network plot of 11 *G. elata* samples shows degree (dot size) and heritability (line color/thickness).

**Figure 4.**
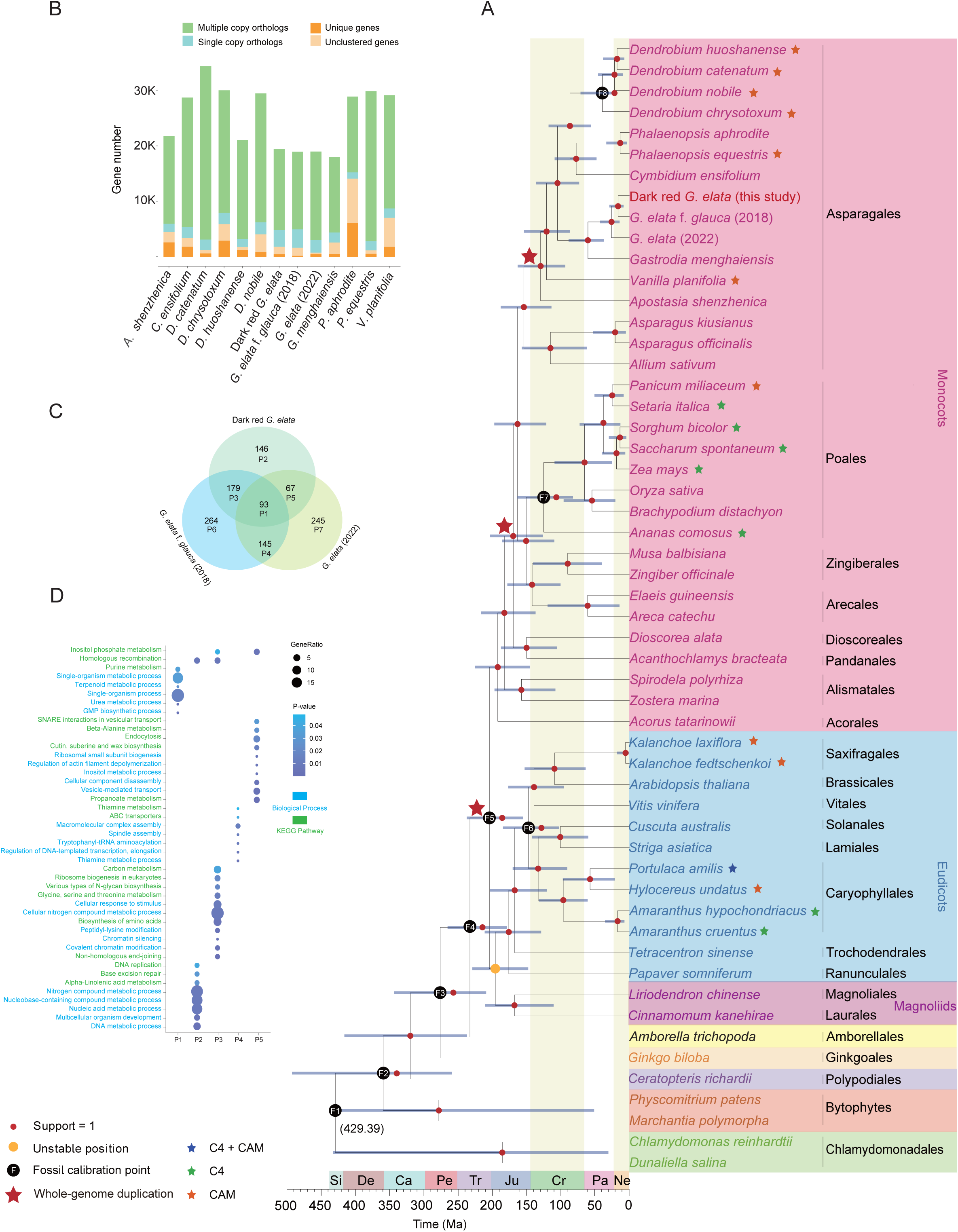
Phylogenetic tree of 54 land plant genomes, with positively selected genes (PSGs) analysis in three *G. elata* forms. A. Summary phylogeny and timescale of 54 land plants. Nodes’ blue bars = 95% credibility intervals of dates. B. Bar graph of protein-coding gene counts in 13 orchids. C. Venn diagram of PSGs in three *G. elata* forms: P1 (common), P3-P5 (pairwise shared), P2, P6, P7 (unique). D. Functional enrichment map of GO and KEGG for common PSGs (P1-P5 refer to C).

**Figure 5.**
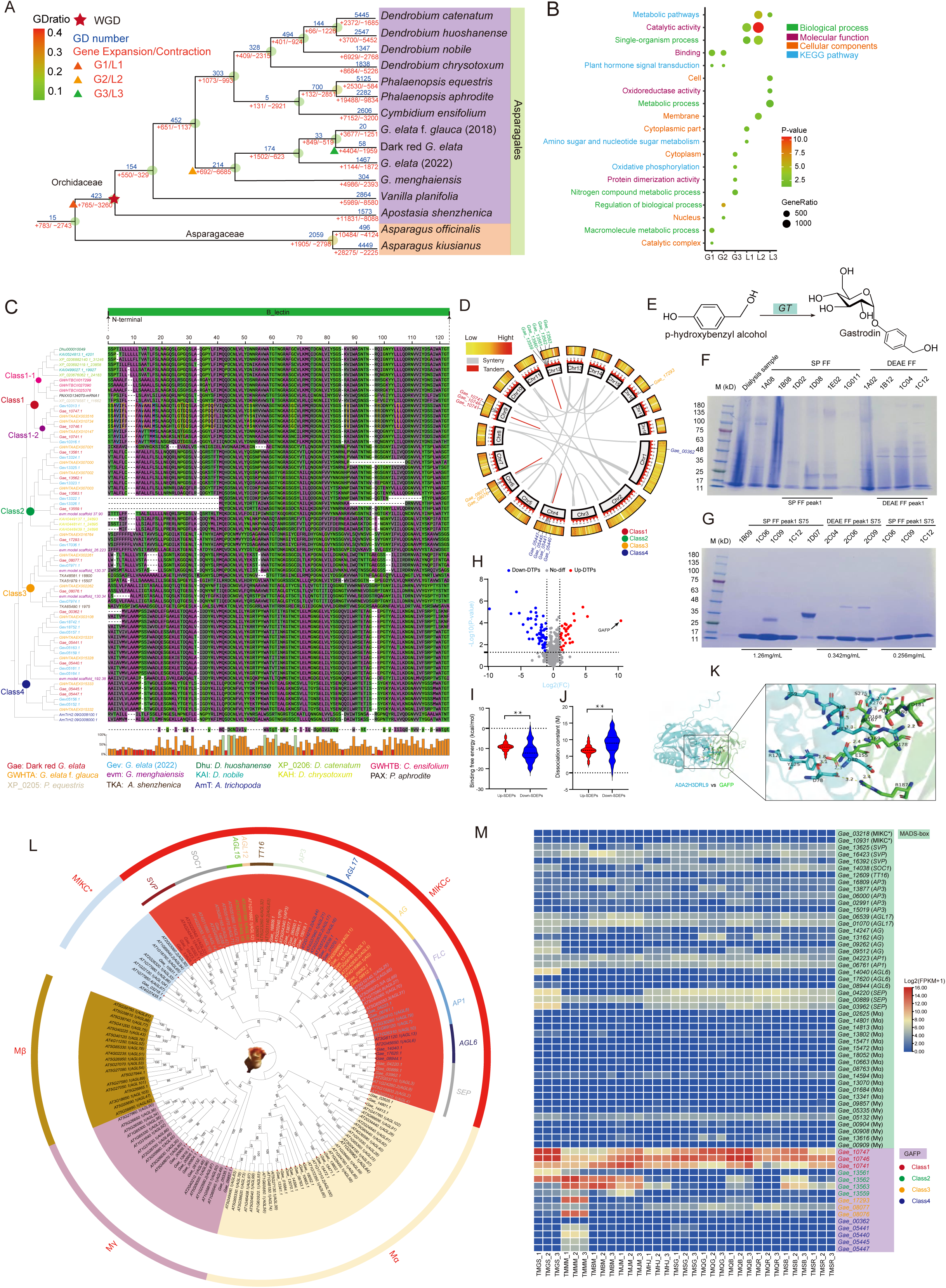
Evolution and functional predictions of gain/loss gene families, the *gastrodia* antifungal protein (GAFP), *MADS-box* gene families in *G. elata*, and the research on the interaction between GAFP and proteins of *A. gallica*. A. The gene family gain/loss and WGD events in 13 orchidaceae plants. B. The KEGG and GO functional enrichment analyses of gain and loss genes in Orchidaceae, *Gastrodia*, and dark-red G*. elata*. G1/L1, G2/L2, and G3/L3 respectively represent the gene families with gain and loss events at the most recent common ancestral (MRCA) node of *Orchidaceae*, the MRCA node of the *Gastrodia*, and dark-red G*. elata*. C. Multiple sequence alignment of 73 GAFP proteins in 13 plant genomes. Unique amino acid residues for Class1-1 highlighted in blue; specificity within Class1 in red; uniqueness to Gea_10746 and Gea_10747 in yellow. Residues with >50% consensus are shown in lowercase, and those with >95% consensus in uppercase. D. The Circos plot of the tandem duplications of the *GAFP* genes in the genome of dark-red G*. elata*. E. Putative gastrodin biosynthetic pathway in *G. elata*. GT, glycosyltransferase. F. The SDS-PAGE analysis of the *G. elata* tuber proteins during the purification stage using Diethylaminoethyl Sepharose Fast Flow (DEAE-SP FF). In the SDS-PAGE diagram, SP FF and DEAE FF refer to the eluates from cation exchange chromatography and anion exchange chromatography, respectively. G. The SDS-PAGE analysis of Elution Peak 1 from DEAE Sepharose FF after purification using Sephadex G-75 gel filtration chromatography. The SP FF peak1 S75 and DEAE FF peak1 S75 refer to the eluates corresponding to the further purified samples of the cation exchange chromatography peak 1 eluate and anion exchange chromatography peak 1 eluate, respectively, using Sephadex G-75 molecular sieve chromatography. H. The volcano plot illustrates significantly differentially enriched proteins. The −log_10_(*P*-value) is plotted against the log_2_(fold change). The non-axial vertical lines denote ±2-fold change while the non-axial horizontal line denotes *P* = 0.05, which denotes the significance threshold (prior to logarithmic transformation). I. The violin plot of free binding energy for significantly differentially enrichment proteins. Red and blue violin plot represent the distribution of free binding energy for proteins with significantly increased abundance and proteins with significantly decreased abundance, respectively. J. The violin plot of dissociation constants for significantly differentially abundant proteins. Red and blue violin plot represent the distribution of dissociation constants for proteins with significantly increased abundance and proteins with significantly decreased abundance, respectively. K. The structural diagram of residue interactions between candidate proteins A0A2H3DRL9 and GAFP. This structural diagram illustrates the molecular interface between the candidate protein A0A2H3DRL9 (in blue) and GAFP (in green), highlighting key interacting residues within 5 Å. Hydrogen bonds are represented as yellow dashed lines, with distances (in Å) and interacting residue sites annotated. L. The *MADS-box* gene family tree in dark-red G*. elata*. The different colors on the outermost ring represent different subfamilies of the *MADS-box*. M. The heatmap of *MADS-box* and *GAFP* gene expression in *G. elata* tissues: capsule (TMGS), immature tuber (TMMM), developing tuber (TMBM), mature tuber (TMJM), flower and stem (TMHJ), and stalk color forms (dark green TMSG, light green TMQG, light brown TMQB, light red TMQR, dark brown TMSB, dark red TMSR).

**Figure 6.**
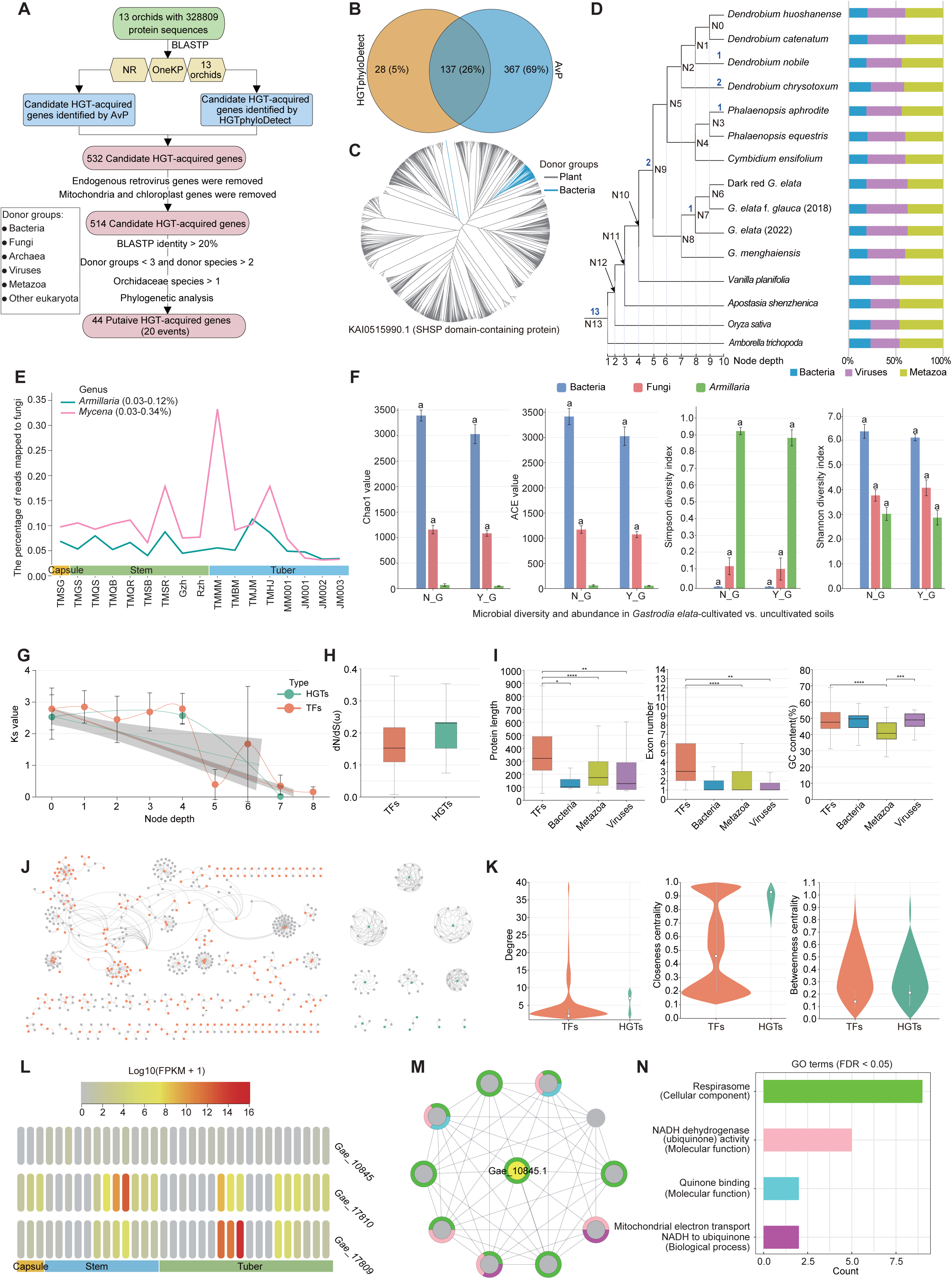
Identification and characteristics of horizontal gene transfer (HGT) in 13 Orchidaceae plants. A: Overview workflow for identifying Orchidaceae candidate HGT-acquired genes. Green oval boxes represent protein sequences, yellow hexagonal boxes indicate BLAST alignment datasets, blue rectangular boxes denote the two software tools used for identifying HGT-acquired genes, and red oval boxes represent the candidate HGT-acquired gene sets obtained at each filtering step. The text on the arrows between each analytical step indicates the corresponding screening process. B: Venn chart showing the number and proportion of candidate HGT-acquired genes identified by HGTphyloDetect and AvP software. C: Molecular phylogeny of *KAI0515990.1* and its homologous genes. Each branch represents a species, with gray branches representing green plants and blue branches representing bacteria. D: Phylogenetic tree of 13 Orchidaceae species and 2 outgroups with origins of HGT-acquired genes. Numbers above branches indicate gene family counts. ‘N’ indicates the node number. Bar charts show the proportions of different donor-mediated HGT events. Vertical gray lines descending from each node demarcate node depth levels. E: Line chart showing the proportion of reads from 17 RNA-seq datasets mapped to five *Armillaria* species and five *Mycena* species. The pink lines represent the genus *Armillaria*, and the green lines represent the genus *Mycena*. F: Bar chart showing microbial diversity and abundance in soil from a controlled experiment comparing plots with (Y_G) and without (N_G) *G. elata* cultivation. Chao1 (Chao1 richness estimator), ACE (Abundance-based Coverage Estimator), Shannon (Shannon diversity index), and Simpson (Simpson diversity index), which are used to estimate the total species richness (Chao1, ACE), community diversity (Shannon), and dominance (Simpson) of the microbial community. G: Line chart illustrating the variation in Ks values of HGT-acquired genes and transcription factor genes across increasing phylogenetic node depths. The two trend lines (green and orange) represent the Ks value trajectories of HGTs and TFs, respectively, with gray shaded areas indicating 95% confidence intervals. Node depth references Figure 6D. H: Boxplot showing the dN/dS (ω) values for HGT-acquired genes and transcription factor genes. I: Boxplot showing a comparison of HGT-acquired genes and transcription factor genes in terms of protein length, exon number, and GC content. J: The protein interaction network of HGT-acquired genes and transcription factor genes. In the network, orange nodes represent transcription factor genes, green nodes represent HGT-acquired genes, and gray nodes represent proteins in the database that interact with either HGT-acquired genes or transcription factor genes. K: Violin plot comparing degree distributions in the protein-protein interaction network between HGT-acquired genes and transcription factor genes. L: Heatmap of HGT-acquired genes expression across *G. elata* stages and color forms. M: The protein interaction network of protein *Gae_10845.1* and the bar plot showing GO enrichment results of this network. Different colored blocks around the nodes in the network correspond to the functional categories in the GO enrichment bar plot (Figure 6N). N: Bar plot showing significantly enriched GO terms (*P* < 0.05) in the biological process, cellular component, molecular function category.

### Intraspecific evolutionary relationships of *G. elata*

To investigate the intraspecific evolutionary relationships of *G. elata*, we constructed a pan-genome comprising 11 *G. elata* genomes and one *G. menghaiensis* genome as an outgroup. Phylogenetic analysis was performed based on single nucleotide polymorphism (SNP) and insertion/deletion (INDEL) datasets (Supplementary Table 17). The SNP and INDEL density plots revealed consistent gene density distribution across all 11 *G. elata* and one *G. menghaiensis* samples (Fig. 3A, 3B), suggesting broadly comparable variation landscapes across accessions. Principal component analysis (PCA) revealed distinct clustering patterns among the analyzed samples (Fig. 3C, 3E). *G. menghaiensis* and *G. elata* f. *flavida* each formed separate and distinct clusters. Interestingly, the remaining *G. elata* samples clustered primarily based on tuber morphology rather than stem coloration: samples with fusiform tubers (including light red, light green, and the 2022 sample) grouped together, whereas samples with oval tubers formed an independent cluster. This morphology-based clustering pattern suggests that tuber shape may represent a more reliable taxonomic criterion for classifying *G. elata* forms than the color-based classification system currently adopted in the *Flora of China*.

Phylogenomic analysis further elucidated the evolutionary relationships within *Gastrodia* (Fig. 3D, 3E). *G. menghaiensis* and *G. elata* were confirmed as sister taxa. Within *G. elata*, f. *flavida* formed a sister clade with varieties possessing fusiform tubers, while varieties with oval tubers comprised an independent monophyletic group. Notably, the SNP-based phylogenetic tree did not resolve a clear evolutionary sequence among the described *G. elata* forms (Supplementary Table 19), further supporting the reliance of tuber morphology over stem color in taxonomic classification.

Given the observed similarities in stem color and tuber shape among certain *G. elata* forms, we hypothesized that some cultivated varieties may have originated through interform hybridization. Hybridization signal analysis supported this hypothesis, suggesting that dark-red G*. elata* potentially resulted from crossing between *G. elata* (2022) (Fig. 3G) and other forms (Fig. 3F and Supplementary Table 18). All 11 *G. elata* samples, collected from cultivated germplasm resources, exhibited complex hybridization signals indicative of historical gene flow (Fig. 3G). Additionally, these samples displayed high protein sequence similarities, with minor variations corresponding to differences in flower and stem coloration (Supplementary Fig. 7). Although extensive artificial hybridization has obscured the intraspecific evolutionary history of cultivated *G. elata*, these genetically diverse germplasm resources nonetheless constitute valuable materials for future breeding programs.

### Phylogeny and divergence time estimation of *G. elata*

To investigate the evolutionary relationships of *G. elata* within a broader phylogenetic context, we constructed a species tree utilizing genomes from 54 terrestrial plant species (Fig. 4A and Supplementary Table 20). The phylogenetic analysis revealed that Orchidaceae constitutes a highly diversified plant family, with *Dendrobium* and *Gastrodia* representing two divergent lineages that have developed distinctly different evolutionary strategies. *Dendrobium* has evolved specialized aerial roots with velamen for survival in arid epiphytic environments, whereas *Gastrodia* has adopted a fully mycoheterotrophic lifestyle through obligate symbiosis with fungi.

Molecular clock analysis revealed the divergence timeline within *Gastrodia* and related taxa (Fig. 4A). *Gastrodia* diverged from *Vanilla planifolia* approximately 120.89 million years ago (Mya). Within the genus, *G. menghaiensis* diverged at approximately 60.57 Mya, followed by *G. elata* (2022) at 26.06 Mya, and subsequently *G. elata* f. *glauca* (2018) and dark-red G*. elata* at 16.54 Mya. Despite the general trend among *Gastrodia* species towards possessing tuberous rhizomes and lacking chlorophyllous leaves, *G. menghaiensis* retains vestigial roots, suggesting it represents a more ancestral lineage. In contrast, *G. elata* has fully adapted to mycoheterotrophy, characterized by complete root loss and an entire vegetative cycle completed in symbiosis with *Armillaria* and *Mycena* fungi.

Synonymous substitution rate (Ks) analysis detected whole-genome duplication (WGD) events across 13 Orchidaceae species, with no recent WGD events observed in *G. menghaiensis* or *G. elata* (Fig. 4A and Supplementary Fig. 8). Statistical analysis revealed that *G. elata* possesses significantly fewer genes compared to other orchids (Fig. 4B), likely attributed to the evolutionary loss of genes associated with root and leaf functions. The absence of recent WGD events combined with substantial gene loss suggests that *G. elata* has undergone genome contraction during its adaptation to obligate mycoheterotrophy.

### Positive selection genes of *G. elata*

Using the ETE Toolkit (ete3), we analyzed 1,837 homologous gene clusters from 13 Orchidaceae species to identify genes under positive selection (Fig. 4C). A total of 485,681, and 550 positively selected genes (PSGs) were identified in dark-red G*. elata*, *G. elata* f. *glauca* (2018), and *G. elata* (2022), respectively. Venn diagram analysis revealed that 93 PSGs (P1) were shared among all three varieties, with an additional 179,145, and 67 PSGs shared pairwise (P3, P4, P5) (Fig. 4C). Gene Ontology (GO) and Kyoto Encyclopedia of Genes and Genomes (KEGG) enrichment analyses indicated that these PSGs are predominantly involved in single-organism processes, nitrogen compound metabolic processes, and purine metabolism (Fig. 4D). These positively selected genes likely enhance material transport and energy metabolism, thereby facilitating nutrient absorption from *Armillaria* and supporting *G. elata* growth in the absence of photosynthetic capability.

### Gene duplications, gene family contraction and expansion across *G. elata*

Whole-genome analysis of gene family evolution was conducted across Orchidaceae species. In the most recent common ancestor (MRCA) of Orchidaceae, 423 gene duplication bursts were identified, consistent with previously reported whole-genome duplication events in the orchid MRCA^26^. Gene family dynamics analysis revealed significant contraction at the common ancestor node of 13 orchid species (Fig. 5A, 5B and Supplementary Tables 21–23). Within the *G. elata* lineage, 623 gene families underwent contraction while 1,502 underwent expansion. GO and KEGG functional analyses demonstrated that expanded genes were enriched in nitrogen metabolism, oxidative phosphorylation, and hormone signal transduction, whereas contracted genes were associated with photosynthesis and root-mediated nutrient uptake. This pattern of selective gene family expansion and contraction reflects the genomic adaptation of *G. elata* to its fully mycoheterotrophic lifestyle during symbiotic evolution.

### Molecular evolution of *GAFP* genes in *G. elata* and their interactions with *Armillaria* protein

*Gastrodia* antifungal protein (GAFP) enhances plant resistance to various pathogens through heterologous expression^27, 28^. However, the molecular evolution of GAFP genes in *G. elata* remains largely unexplored. To address this, we constructed a phylogenetic tree using antifungal protein sequences from 13 orchid species, including three *G. elata* forms and *G. menghaiensis* (Fig. 5C and Supplementary Fig. 9). The phylogenetic analysis clustered antifungal protein sequences into four classes (Class 1-4). Notably, *G. menghaiensis* lacked GAFP gene copies in Class 1 but retained copies in Classes 2-4. Multiple sequence alignment revealed distinct differences in four protein domains between Class 1 GAFP sequences and those from Classes 2-4 (Fig. 5C), suggesting recent evolutionary divergence in this gene cluster. Examination of duplication events revealed that GAFP gene family members are distributed across six chromosomes in dark-red G*. elata*: Class 1 on Chr9, Class 2 on Chr12, Class 3 on Chr6 and Chr16, and Class 4 on Chr1 and Chr4 (Fig. 5D). Collinearity analysis demonstrated that tandem duplication represents the primary expansion mechanism for the GAFP family. Among 15 GAFP genes, 11 exhibited four tandem repeat relationships: *Gae_05440*/*Gae_05441*, *Gae_08076*/*Gae_08077*, *Gae_10746*/*Gae_10747*, and *Gae_13559*/*Gae_13560*/*Gae_13561*/*Gae_13562*/*Gae_13563*. These findings indicate that *G. menghaiensis* exhibits a lower molecular evolutionary rate than *G. elata*, retaining vestigial root systems on its tubers, whereas *G. elata* has completely lost root systems and relies entirely on symbiosis with *Armillaria* and *Mycena* for nutrition. Transcriptome analysis of six *G. elata* color forms and five developmental stages revealed progressively increasing GAFP expression from Class 4 to Class 1, with highest expression in Class 1 (Fig. 5I). This pattern suggests that continuous evolution of GAFP structural domains may enhance antifungal capability, enabling *G. elata* to obtain nutrition from fungal symbionts while preventing excessive fungal growth. Among developmental stages, GAFP expression was highest in capsules, followed by immature tuber, developing tuber, and mature tuber (Fig. 5F). High GAFP expression in capsules and immature tubers likely protects seeds from pathogenic fungi (excluding beneficial *Mycena* for germination) and maintains normal tuber development.

To investigate the interaction between *G. elata* and *Armillaria gallica*, GAFP protein (GWHTBDNU009330) was purified from fresh *Gastrodia* tubers and used as bait in biotin pull-down assays^4, 29^. LC-MS/MS analysis identified 26 significantly upregulated *A. gallica* proteins that potentially interact with GAFP^30, 31^. These proteins exhibited diverse physicochemical properties, ranging from 60 to 1,279 amino acids in length, with most having isoelectric points between 5.5 and 7.5 (Supplementary Table 24). The majority (25/26) displayed negative hydrophobicity values, indicating hydrophilic nature suitable for cellular environments^32^. Domain prediction and KEGG pathway analysis revealed enrichment in metabolism, enzyme activity, cellular signaling, and energy production, with significant associations to carbon metabolism and glyoxylate/dicarboxylate metabolism pathways.

Binding capacity between candidate proteins and GAFP was assessed using sequence-based free binding energy prediction (PPA_Pred) and molecular docking analysis. Twenty proteins exhibited absolute free binding energies of 6–13 kcal/mol and −log₁₀ dissociation constants of 5-9 M, indicating significant GAFP binding capacity. Binding parameters differed significantly between upregulated and downregulated proteins (*P* < 0.01) (Fig. 5H, 5I and Supplementary Table 25). Molecular docking evaluation across three dimensions—Docking Score, Confidence Score (>0.7 indicates high binding likelihood), and Ligand RMSD^33^, identified A0A2H3DRL9 as the top candidate with a docking score of −290.73, Confidence Score of 0.9435, and Ligand RMSD of 37.32 Å. The docking interface revealed 13 hydrogen bonds (1.8–3.5 Å) forming a stable interaction network (Fig. 5J)^34^. Proteins A0A2H3DFK3, A0A2H3DQC5, and A0A2H3E9J7 also demonstrated favorable docking interactions (Tables S25-27 and Supplementary Fig.15). These four proteins represent priority candidates for experimental validation of GAFP interactions.

Gastrodin (GAS), a major bioactive compound in dried *G. elata* tubers, combined with para-hydroxybenzyl alcohol (p-HBA) typically exceeds 0.25% content^35^. The final step in GAS biosynthesis involves glycosyltransferase (GT)-mediated conversion of p-HBA to GAS^36, 37^. The *AsUGT* enzyme from *Rauvolfia serpentina* demonstrates superior p-HBA to GAS conversion efficiency in *Saccharomyces cerevisiae* compared to UGT73B6^36^. Using *AsUGT* sequences as queries, BLASTP analysis against 54 land plant genomes identified multiple homologs (Fig. 5E and Supplementary Fig.10). In *G. elata*, 54 *GT* gene copies were identified as *AsUGT* homologs, with three genes (*Gae_12521*, *Gae_11966*, and *Gae_08743*) highly expressed in immature, developing, and mature tuber stages (Supplementary Table 28). This expression pattern corresponds with GAS accumulation during tuber maturation and steaming processes, confirming that GAS biosynthesis occurs primarily within *G. elata*. These findings provide theoretical foundation and technical support for artificial gastrodin synthesis.

### Molecular evolution of the *MADS-box* gene family in *G. elata*

*MADS-box* genes play crucial roles in regulating plant floral organ development, morphology, and flowering time^38^. To investigate their potential involvement in *G. elata*’s unique morphological traits, we analyzed both type I (Mα, Mβ, Mγ subfamilies) and type II (MIKC^C^ and MIKC* subfamilies) MADS-box gene families using *Arabidopsis thaliana* as an outgroup (Fig. 5L). Within type I *MADS-box* genes, the Mα subfamily contained 13 homologs in *G. elata*. However, only *Gae_14594* (FPKM ∼4.60-6.32) and *Gae_13802* (FPKM ∼0.70-1.25) showed detectable expression in flowers and stems, while the remaining homologs exhibited minimal or no expression (Fig. 5I). In the Mγ subfamily, seven homologs were identified, with only *Gae_05132* (FPKM ∼8.20-14.00) weakly expressed across five developmental stages and six stem color forms (Fig. 5M). Notably, no Mβ subfamily homologs were identified in *G. elata*. In *Arabidopsis*, certain Mα, Mβ, and Mγ genes (e.g., *AGL34, AGL35, AGL36, AGL37, AGL38, AGL62, AGL80, AGL86, AGL89, AGL90*) are associated with endosperm development and proliferation^39^. The absence of Mβ genes in *G. elata* may reflect an evolutionary adaptation correlated with the reduced endosperm characteristic of its seeds. Given that *G. elata* produces numerous tiny seeds that germinate upon encountering mycorrhizal fungi (e.g., *Mycena*) in leaf litter, relaxed selection pressure on endosperm formation appears plausible. Within type II *MADS-box* genes, 26 homologs were identified with MIKC^C being predominant. These genes were examined in the context of the ABCDE model of floral organ development established in *Arabidopsis*, where different *MADS-box* gene combinations govern the identity of sepals, petals, stamens, carpels, and ovules. In *G. elata*, class A homologs (AP1-like: *Gae_04223, Gae_06761*), class B homologs (AGL17-like: *Gae_06539, Gae_01070*), and class E homologs (SVP-like: *Gae_16423*; SEP-like: *Gae_00889*) were expressed across multiple developmental stages and stem color variants (Fig. 5M). Since *Arabidopsis* A, B, and E genes influence sepal and petal formation^40^, their *G. elata* counterparts may contribute to the fusion of these organs into a perianth tube. Two SVP-like genes (*Gae_16423* and *Gae_13625*) exhibited higher expression levels, potentially involved in maintaining fused floral structures. In contrast, several AP3-like B-class genes and C/D-class genes (AG and AGL11 homologs) were present but showed low or negligible expression. *G. elata* also lacks certain MADS-box homologs present in *Arabidopsis*, including FLC (associated with vernalization response^31, 32^, AGL15 (endosperm development^41^), AGL12 (root formation^42^), and AGL6 subfamilies (flowering induction^43^). The absence or reduced expression of these genes correlates with *G. elata*’s distinctive life history characteristics: lacking roots and leaves, flowering during relatively warm seasons, and producing seeds with highly reduced endosperm. These findings suggest that *MADS-box* gene family evolution has contributed to the unique morphological adaptations of *G. elata* to its mycoheterotrophy lifestyle.

### Degeneration of photosynthesis function in *G. elata*

To elucidate the molecular basis underlying the loss of photosynthesis in *G. elata*, we systematically examined genes involved in carbon fixation and chlorophyll biosynthesis. Two phosphoenolpyruvate carboxylase (*PPC*) genes, Gae_14940 (*PPC1M1*) and Gae_12186 (*PPC1M2*), were identified in the *G. elata* genome and clustered phylogenetically with *PPC* homologs from C₃/CAM plants (Supplementary Fig. 11). However, both genes displayed extremely weak and spatially restricted expression, with *PPC1M1* detected only in light-red stems and *PPC1M2* confined to light-green stems, while transcription was nearly undetectable in stems of other colours and completely absent in tubers (Supplementary Fig. 12), indicating minimal functional contribution to carbon assimilation. Strikingly, no nuclear-encoded Rubisco small subunit gene (*rbcS*) was identified in *G. elata*. Given the essential role of *rbcS* in Rubisco holoenzyme assembly, its complete loss precludes functional Calvin–Benson cycle activity, providing direct molecular evidence for the irreversible loss of autotrophic carbon fixation in this species. This *rbcS*-centered degeneration represents a decisive step in dismantling photosynthetic capacity during the transition to obligate mycoheterotrophy.

Analysis of the chlorophyll biosynthesis pathway identified 14 enzyme-encoding genes retained in the *G. elata* genome. Among these, chlorophyllide a oxygenase (CAO; Gae_07292), which catalyzes the final conversion of chlorophyll a to chlorophyll b, exhibited relatively high expression levels, whereas the remaining pathway genes showed uniformly low transcriptional activity (Supplementary Fig. 13). This expression pattern suggests selective retention of individual chlorophyll-related components rather than maintenance of a functional photosynthetic apparatus. Pan-genome analysis further revealed a non-random distribution of photosynthesis-related genes between core and non-core gene sets. While core genes showed weak and statistically non-significant enrichment for photosynthesis-associated Gene Ontology (GO) terms, non-core genes were significantly enriched for photosynthesis-related pathways (Supplementary Table 15), indicating relaxed selective constraints and lineage-specific retention. Collectively, these results demonstrate that *G. elata* has undergone advanced reductive evolution of the photosynthetic machinery, driven by the loss of the nuclear *rbcS* gene, while retaining a subset of photosynthesis-related genes that are likely repurposed for alternative roles. These retained genes may contribute to light-responsive regulation, photomorphogenesis, circadian rhythm control, or flowering-time regulation, reflecting a functional shift from autotrophy to complete mycoheterotrophy.

### Systematic identification and evolutionary analysis of horizontal gene transfer events in Orchidaceae plants

To systematically investigate HGT events and their evolutionary characteristics in 13 orchid species (Supplementary Table 29), we employed a rigorous three-step identification pipeline (Fig. 6A), including initial screening of candidate genes (Fig. 6B, Supplementary Table 30, S31), multi-dimensional filtering, and phylogenetic validation (Fig. 6C). In these 13 orchid species, we identified 44 horizontally transferred genes belonging to 20 distinct gene families (Fig. 6D, Supplementary Table 32). Notably, although *G. elata* maintains a long-term nutritional symbiotic relationship with *Armillaria*/*Mycena* fungi—a relationship generally considered to create favorable conditions for horizontal gene transfer due to their prolonged close coexistence and potentially resulting in a higher frequency of HGT events—our analysis detected no HGT events at the ancestral node of *G. elata* (node N8 in Fig. 6D). Instead, we detected only one viral-derived HGT event at the common ancestor node of *G. elata* f. *glauca* (2018) and *G. elata* (2022) (node N7 in Fig. 6E), involving two genes (*Gev01277.1* in *G. elata* (2022) and *GWHTAAEX018452* in *G. elata* f. *glauca* (2018)). To further validate this finding, we conducted additional experiments: (1) Transcriptome analysis showed that the proportion of RNA-seq reads from 17 *G. elata* transcriptomes mapped to fungal genomes was extremely low (0.03%-0.34%) (Fig. 6E, Supplementary Table 33), indicating that fungal genes are rarely actively expressed in *G. elata* tissues. (2) Rhizosphere microbial diversity analysis revealed no significant differences in α-diversity (including Chao1 species richness, ACE coverage, Shannon diversity, and Simpson dominance indices) of the microbial communities in the rhizosphere soil, regardless of whether *G. elata* was cultivated (p > 0.05). More importantly, no *Mycena* fungi were detected in any of the samples (Fig. 6F). These results further demonstrate that no HGT events occurred between *G. elata* and its symbiotic fungi, suggesting that their symbiotic relationship is more likely maintained through metabolic interactions.

Based on the identified HGT genes, we further analyzed their molecular evolutionary characteristics. Molecular evolution analysis showed that the synonymous substitution rate (Ks) of HGT genes decreased with evolutionary depth and was significantly lower than that of the orchid’s own transcription factor (TF) genes (Fig. 6G, Supplementary Table 34). This suggests that these HGT genes may be relatively “young.” Meanwhile, HGT genes exhibited higher ω values (dN/dS, the ratio of nonsynonymous to synonymous substitution rates), indicating positive selection and potential adaptive evolution. In contrast, TF genes exhibited relatively low *dN*/*dS* (ω) values, indicating their evolutionary conservation and suggesting that they are primarily under purifying selection (Fig. 6H). Additionally, HGT genes generally had shorter protein sequences, fewer exons, and lower GC content (Fig. 6I, Supplementary Table 35), reflecting their early evolutionary state before full optimization.

To understand the expression patterns and functions of HGT genes in their host species, we performed protein-protein interaction (PPI) network analysis. The results showed that the overall complexity of the PPI network formed by HGT genes was significantly lower than that of TF gene networks (Fig. 6J-K, Supplementary Fig.15). However, in-depth analysis of individual gene network attributes revealed a seemingly paradoxical phenomenon: HGT genes exhibited higher connectivity, closeness centrality, and betweenness centrality (Fig. 6K). We speculate that this apparent contradiction arises from hierarchical differences in network architecture: while TF genes have broader interaction capabilities, their interactions are dispersed across multiple functional modules, presenting a more distributed linkage. In contrast, the interaction network of HGT genes exhibits distinct localization features. Additionally, the significant numerical disparity between TF genes and HGT genes (with TF genes far outnumbering HGT genes) may also influence the comparison of these metrics to some extent. Furthermore, we observed that among the HGT genes in *G. elata*, three genes (Fig. 6L, Supplementary Table 36) were expressed in *G. elata*. One of these, *Gae_10845*, had 10 directly interacting proteins (Fig. 6M). Gene Ontology (GO) enrichment analysis indicated that this gene and its interaction partners are involved in energy metabolism-related processes in the mitochondrial inner membrane (Fig. 6N), suggesting a potential role in host energy metabolism.

In summary, this study identified 20 HGT events in orchids. Despite the long-term symbiotic relationship between *G. elata* and fungi, no fungal-derived HGTs were detected in its genome, with only virus-derived transfer events observed. This indicates that the symbiotic relationship primarily relies on metabolic interactions. The recent origin (low Ks), positive selection (high ω), and non-optimized sequence features (short CDS, low GC content) of HGT genes contrast sharply with the conservation of TF genes. Their roles in basic metabolic regulation (e.g., energy metabolism) provide new insights into orchid-microbe coevolution.

### Transporter gene family analyses provide insights into mutualism in *G. elata*

A. *G. elata* is an obligately mycoheterotrophic orchid characterized by non-endospermic seeds and the complete absence of roots, rhizomes, and leaves. Our analysis confirmed that *G. elata* lacks Rubisco, indicating an inability to acquire nutrients via photosynthesis. Instead, *G. elata* depends entirely on symbiosis with *Armillaria mellea* for carbon (C) and nitrogen (N) acquisition, where the fungus decomposes organic matter and transfers derived nutrients to *G. elata* during colonization^44, 45^.

Based on the dark-red G*. elata* genome, we identified 95 sugar-related genes encoding monosaccharide transporters, sucrose synthases, trehalose, and related enzymes. These genes exhibited tissue-specific expression patterns, suggesting that *G. elata* acquires glucose, fructose, galactose, sucrose, and trehalose from fungi (Fig. 7, Figures S17-S32 and Tables S37-39). Notably, trehalose from fungi represents a major carbon source for *G. elata*. Increased trehalase gene copy numbers in *G. menghaiensis* indicate the capacity to decompose fungal trehalose into glucose^11, 46, 47^. Phylogenetic analysis revealed that *Gastrodia* trehalose differ significantly from those in other orchids, monocots, and dicots (Fig. 7C and Supplementary Table 38), further confirming adaptation to mycoheterotrophy through trehalose-derived glucose utilization. The sugar transporter protein (STP) family exhibits marked expansion in dark-red G*. elata*, with 18 copies identified (Supplementary Table 38). Among these, *Gae_05439*, *Gae_10698*, and *Gae_15520* are orthologous to *Arabidopsis* AtSTP3, which mediates energy-dependent transport of hexoses and pentoses^48, 49^, and exhibit elevated expression in mature tubers (Fig. 7B and Supplementary Table 39). *Gae_07301*, orthologous to AtSTP14 which exhibits strict galactose transport specificity^50^ , was identified as a positively selected gene (Supplementary Fig.51) and shows high expression in immature tubers and capsules. The vacuolar glucose transporter (VGT) gene *Gae_08349*, whose *Arabidopsis* ortholog mediates glucose sequestration into vacuoles^51^ , is specifically expressed in flowers and stems. The presence of sucrose synthase (SUS), invertase (Inv), and sucrose transporter (SUT) genes indicates the capacity for direct utilization of fungal sucrose or its hydrolysis products^20, 52, 53^. Furthermore, genes encoding maltase, maltose transporters, and starch hydrolases enable decomposition of fungal polysaccharides into absorbable monosaccharides. Comparative analysis across *Gastrodia* species, orchids, monocots, and dicots revealed that obligately mycoheterotrophic *Gastrodia* species preferentially acquire glucose via fungal trehalose hydrolysis, with efficient transport mediated by STP and VGT family members, supplemented by direct sucrose uptake and polysaccharide degradation pathways (Fig. 7C and Supplementary Table 38).

**Figure 7.**
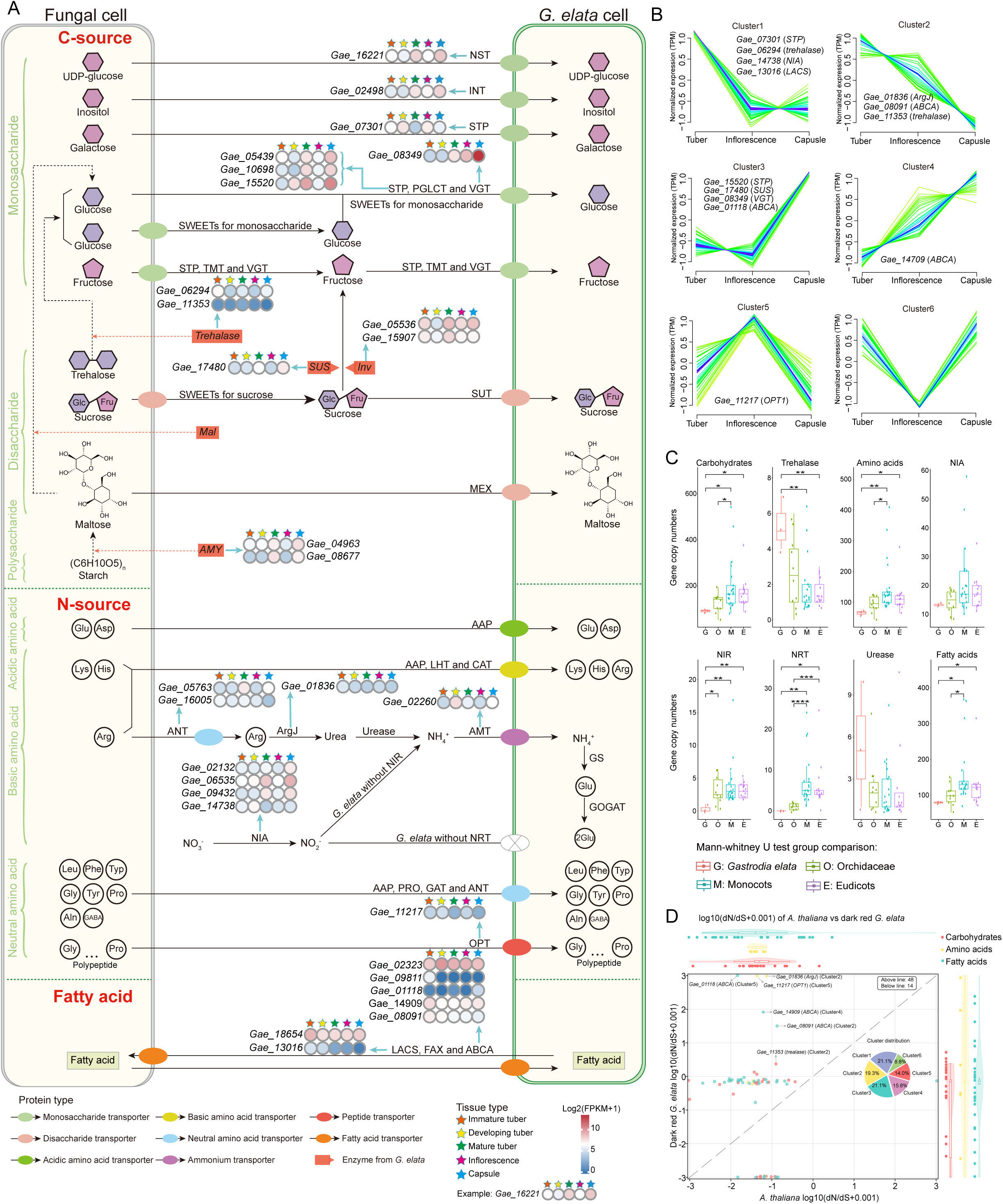
Identification of key enzymes in sugar, amino acid, fatty acid, and gastrodin metabolic pathways, and diagram of homologous copy gene expression levels in *G. elata*. This diagram consists of four core sections, as detailed below: A. (a) C-source. The left side represents the fungal cell wall, covering three types of carbohydrates: monosaccharides, disaccharides, and polysaccharides; the right side represents the *G. elata* cell wall, marking the types of carbohydrates transported from fungi to *G. elata*. Monosaccharide transporters, sucrose transporters, maltose transporters, and sugar metabolism enzyme genes are clearly labeled in the diagram. Sugar-related abbreviations (including sugar transporters, sugar metabolic enzymes, and sugar molecules): UDP-Glc (uridine diphosphate glucose), Ino (inositol), Gla (Galactose), Glc (glucose), Fru (fructose), NST (neutral sugar transporter), INT (inward transporter), STP (sugar transport protein, or a member of the monosaccharide transporter family), PGLCT (putative glucose-lactose transporter), VGT (vacuolar glucose transporter), TMT (tonoplast monosaccharide transporter), *SUS* (sucrose synthase), Inv (invertase), SUT (sucrose transporter), *AMY* (amylase), *Mal* (maltose), *MEX* (maltose exporter). (b) N-source. The left side shows the types of amino acids present in fungi, including acidic amino acids, basic amino acids, neutral amino acids, and polypeptides; the right side shows the types of amino acids that *G. elata* may acquire from fungi via transport. Abbreviations for amino acids and nitrogen-containing compounds (including amino acid transporters and nitrogen metabolism enzymes): Standard amino acids: Glu (glutamic acid), Asp (aspartic acid), Lys (lysine), His (histidine), Arg (arginine), Phe (phenylalanine), Gly (glycine), Pro (proline), Trp (tryptophan), Ala (alanine), Tyr (tyrosine), Leu (leucine); Non-standard nitrogen-containing metabolites: GABA (γ-aminobutyric acid, involved in nitrogen metabolism); Transport/metabolism-related proteins: AAP (amino acid permease), LHT (lysine/histidine transporter), CAT (cationic amino acid transporter), ANT (amino acid-nitrogen transporter), *ArgJ* (ornithine acetyltransferase), AMT (ammonium transporter), *GS* (glutamine synthetase), *GLU* (glutamate dehydrogenase), *NIR* (nitrite reductase), *NIA* (nitrate reductase), NRT (nitrate transporter), PRO (proline transporter), GAT (GABA transporter), OPT (oligopeptide transporter), PTR (peptide transporter). (c) Fatty acid transport. The left side shows fatty acids in fungi, and the right side shows fatty acids in *G. elata*. Fatty Acid-related abbreviations: LACS (long-chain acyl-CoA synthetase), FAX (fatty acid exporter), ABCA (atp-binding cassette subfamily A). Different types of transporters in the diagram are distinguished by different colors, and the expression levels of some homologous genes in *G. elata* are displayed in the diagram; different colored pentagrams represent different developmental stages of *G. elata*, including immature tuber, developing tuber, mature tuber, flower and stem, and capsule. B. Time-series profiles of expression levels of gene copies associated with energy metabolism in dark-red G*. elata*. Through clustering analysis, these genes are divided into 6 expression pattern clusters (Cluster 1 to Cluster 6), with each subplot corresponding to the expression trend of genes in a specific cluster across different tissues. The x-axis represents 3 tissue types, in sequence: Tuber: including tuber tissues at three developmental stages, immature tuber, developing tuber, and mature tuber; Inflorescence: inflorescence tissues; Capsule: capsule tissues. The y-axis indicates the degree of change in gene expression levels relative to the baseline, ranging from −1 to 1. The sign (positive/negative) and magnitude of the values reflect differences in the expression levels of genes in the corresponding tissues. Each curve in each cluster represents the expression trend of a single gene. C. The Mann–Whitney U-tests represent comparisons of the copy numbers of relevant genes among the four plant groups. Abbreviations of the four plant groups: G (*G. elata*), O (Orchidaceae excluding *G. elata*), M (Monocots excluding Orchidaceae), E (Eudicots). Significance symbols for Mann–Whitney U-test results: *** indicates *P* < 0.001, ** indicates *P* < 0.01, and * indicates *P* < 0.05. D. Comparison of log_10_(dN/dS + 0.001) ratios of genes related to carbohydrates, amino acids, and fatty acids between *A. thaliana* and dark-red G*. elata*. The scatter plot shows the correlation analysis of gene dN/dS values between *A. thaliana* and dark-red G*. elata* (the red dashed line represents the diagonal, indicating equal dN/dS values between the two species). The box plots at the top and right present the distribution characteristics of dN/dS values of genes related to the three types of metabolites, where blue represents carbohydrates, orange represents amino acids, and green represents fatty acids.

We identified 67 nitrogen metabolism-related genes in *G. elata*, including those encoding amino acid-polyamine–choline (APC) transporters^54^, neutral amino acid transporters (ANT)^55^, nitrate reductase (NIA1)^56^, nitrite reductase (NIR1)^57^, ArgJ enzyme^58^, NRT enzyme^44^, AMT enzyme^59^, urease^60^, and oligopeptide transporters (Fig. 7A, Figures S33-47 and Tables S37-38). Notably, NIR and NRT genes were absent, indicating *G. elata* cannot acquire environmental nitrate as a nitrogen source, consistent with *G. menghaiensis*^20^. However, *G. elata* retains multiple NIA gene copies (*Gae_02132*, *Gae_06535*, *Gae_09432*, and *Gae_14738*), expressed across all developmental stages with moderately elevated levels in aerial tissues (Fig. 7A and Supplementary Fig.51). Among these, *Gae_09432* and *Gae_14738* exhibit signatures of positive selection. The retained AAT, ANT, AMT, ArgJ, and urease genes collectively support organic nitrogen and ammonium uptake^61^ (Supplementary Table 37, 38). The ANT gene *Gae_06393* is highly expressed in capsules, meeting nitrogen demands for floral stem growth and fruit development. Arginine transported by ANT is hydrolyzed to urea by arginase, then decomposed to ammonium by urease, and absorbed via AMT^55^. Glutamine synthetase enables assimilation of transported ammonium into amino acids^61^. Complete oligopeptide transporter (OPT) genes allow direct uptake of macromolecular peptides from symbiotic fungi, expanding organic nitrogen utilization^62^.

The *G. elata* genome retains a complete triacylglycerol (TAG) synthesis pathway, including acetyl-CoA carboxylase (Accase), β-ketoacyl-ACP synthase (KAS), fatty acyltransferase (FAT), and diacylglycerol acyltransferase (DGAT)^63^ (Supplementary Table 38). This pathway integrity indicates independent fatty acid synthesis capability. FAT gene expression was detected alongside various long-chain fatty acids at different developmental stages (Supplementary Table 40), with linoleic acid providing energy during tuber growth and seed germination. Multiple transporter families facilitate fatty acid transport and storage. FAX family transporters transport long-chain fatty acids from plastids to the endoplasmic reticulum, with *Gae_02829* expressed in immature and developing tubers^64^ (Supplementary Fig.48 and Tables S37-38). ABCA family transporter *Gae_02323* is highly expressed at all developmental stages, while *Gae_09811* shows significantly higher expression in immature tubers^65^ (Supplementary Fig.50 and Supplementary Table 39), potentially transporting fatty acids into tuber cells for storage. Additionally, *Gae_18654* and *Gae_13016*, homologous to *AtLACS3*, show higher expression in immature tubers (Supplementary Fig.49 and Supplementary Table 39), potentially assisting FAX and ABCA transporters in fatty acid transmembrane transport^66, 67^

As an obligate mycoheterotroph, *G. elata* has developed unique evolutionary strategies through long-term symbiosis with *A. mellea*. Energy metabolism-related genes have undergone targeted adaptive adjustments: numerous genes involved in autonomous energy synthesis have been lost, with core gene copy numbers much lower than in green plants (Fig. 7C), reducing genetic load and energy consumption. Conversely, retained key genes including trehalase, STP family genes, and ArgJ exhibit significant positive selection with elevated dN/dS ratios compared to *Arabidopsis* (Fig. 7D and Supplementary Fig.51), enabling more efficient utilization of fungal nutrients. However, subcellular localization of these nutrient transport-related proteins remains unclear and requires further experimental verification.

## Discussion

*Gastrodia elata* is a fully mycoheterotrophic orchid that has lost functional photosynthesis and relies on fungal partners for carbon and nutrients, making it an exceptional system for investigating reductive genome evolution, symbiotic adaptation, and the evolutionary limits of plant–fungus interactions. By generating the first chromosome-level genome assembly of a dark-red *G. elata* accession and constructing a pan-genome encompassing 12 *Gastrodia* genomes, we provide a comprehensive genomic framework for understanding the evolutionary transition from autotrophy to obligate mycoheterotrophy. Our analyses challenge several prevailing assumptions regarding taxonomy, horizontal gene transfer, and symbiotic integration, while revealing coordinated patterns of gene loss, functional repurposing, and adaptive innovation.

A prominent feature emerging from our pan-genome analysis is the pervasive hybridization detected across all examined *G. elata* genomes. The widespread admixture signals observed among cultivated accessions suggest that extensive hybridization has occurred during the history of artificial propagation, likely exacerbated by insufficient control of parental lineages. This finding has important implications for interpreting intraspecific diversity in this economically important medicinal plant. Notably, phylogenetic reconstruction based on genome-wide SNPs indicates that tuber morphology provides a more reliable indicator of genetic relationships than stem color, which has traditionally served as the primary taxonomic criterion in the *Flora of China*. These results call for a re-evaluation of color-based classification schemes and highlight the necessity of using germplasm with well-documented provenance in future population genetic and breeding studies to avoid confounding effects of anthropogenic hybridization.

A defining hallmark of obligate mycoheterotrophy is the irreversible dismantling of the photosynthetic machinery. In *G. elata*, this transition is marked by the complete loss of the nuclear-encoded Rubisco small subunit gene (*rbcS*), a key component required for Rubisco holoenzyme assembly and Calvin-Benson cycle activity. Loss of *rbcS* alone is sufficient to abolish functional carbon fixation, regardless of residual plastid gene content, and therefore represents a decisive nuclear-encoded molecular “point of no return” in the evolution toward full heterotrophy. In contrast to mixotrophic orchids that retain partial photosynthetic capacity, *G. elata* exhibits advanced reductive evolution of the photosynthetic apparatus, consistent with its strict dependence on fungal carbon sources. Intriguingly, a subset of photosynthesis-associated genes, including components of the chlorophyll biosynthesis pathway, is retained and transcriptionally active. Rather than supporting photosynthesis, these genes are likely maintained under alternative selective constraints and may have been co-opted for roles in light-responsive regulation, photomorphogenesis, circadian rhythm, or developmental timing. Such functional repurposing underscores that reductive genome evolution is not a passive process of gene loss, but involves selective retention and reorganization of ancestral pathways.

Despite prolonged and intimate associations with fungal partners, we detected no evidence of fungal-derived horizontal gene transfer (HGT) in *G. elata*. This absence is striking given that theoretical models often predict increased opportunities for genetic exchange under sustained physical intimacy. Our conclusion is supported by both phylogenomic analyses and transcriptomic data, which revealed extremely low proportions of fungal-derived reads in plant tissues. These findings contrast with well-documented cases of adaptive HGT in other plant-microbe interactions, such as the acquisition of Fhb7 in wheat-*Fusarium* systems^15^. Together, our results indicate that intimate symbiosis alone is insufficient to drive genetic integration and suggest that *G. elata*-fungus mutualism is maintained through metabolic and regulatory coordination rather than genetic fusion. This challenges the assumption that symbiosis necessarily promotes HGT and contributes to a broader understanding of the conditions under which inter-kingdom gene transfer is evolutionarily favored. Possible explanations include physical barriers to DNA transfer across plant-fungus interfaces and the rapid purging of maladaptive foreign genes by purifying selection.

Maintenance of a stable symbiosis in the absence of genetic integration likely requires precise molecular control of fungal growth. In this context, the *Gastrodia* antifungal protein (*GAFP*) gene family emerges as a key evolutionary innovation^12, 27, 28^. Our phylogenetic analyses identify four distinct GAFP clades with divergent evolutionary trajectories, indicating repeated functional diversification. Notably, Class 1 GAFP exhibits modified domain architecture, elevated expression across developmental stages, and is absent from the early-diverging species *G. menghaiensis*^11^, suggesting that it represents a relatively recent innovation in *G. elata*. We propose that structural modifications in Class 1 GAFP enhance its ability to suppress excessive fungal proliferation while preserving mutualistic interactions. This hypothesis is supported by previous observations that GAFP accumulates at high levels in immature tubers, a developmental stage at which protection against fungal overgrowth is likely critical^68^. Consistent with this hypothesis, biotin-labelled GAFP pull-down assays identified multiple *Armillaria* proteins with potential binding capacity, providing a molecular entry point for dissecting host-mediated regulation of fungal partners. However, the mechanisms by which GAFP discriminates between beneficial symbiosis and pathogenic overgrowth remain to be elucidated.

Our integrated genomic and transcriptomic analyses reveal that *G. elata* has evolved a comprehensive nutrient exchange system that is more elaborate than previously appreciated^11, 20, 55^. The complete loss of photosynthetic capacity necessitates highly efficient nutrient acquisition strategies. Our integrative analyses reveal that *G. elata* has evolved a sophisticated and actively optimized nutrient exchange framework. Carbon acquisition is primarily mediated through glucose derived from fungal trehalose hydrolysis, supported by expansion of trehalase-encoding genes^11, 69^ and a markedly enlarged repertoire of sugar transporter proteins (STPs). Additional pathways, including sucrose uptake via SUT transporters and polysaccharide degradation by diverse hydrolases^20^, further enhance metabolic flexibility. In parallel, *G. elata* has abandoned nitrate uptake and reduction, yet retains a complete organic nitrogen assimilation system. Neutral amino acid transporters, together with oligopeptide transporters of the OPT and PTR families^62^, facilitate efficient acquisition of fungal-derived nitrogen. Importantly, several key genes involved in nutrient acquisition, including *trehalase*, *STPs*, and *ArgJ*, show signatures of positive selection, indicating ongoing adaptive optimization rather than passive reliance on fungal partners. These findings highlight that obligate mycoheterotrophy involves not only extensive gene loss, but also active molecular adaptation of retained metabolic pathways.

Several limitations of this study warrant consideration. The historical origins and evolutionary consequences of extensive hybridization among cultivated *G. elata* remain incompletely resolved. In addition, the protein-protein interactions inferred between GAFP and fungal targets require direct functional validation, and the precise spatial localization of nutrient transporters at symbiotic interfaces remains to be determined. Future studies integrating controlled breeding experiments, cellular localization analyses, and functional assays will be essential to translate genomic inferences into mechanistic understanding. Comparative analyses across independent mycoheterotrophic lineages will further clarify the generality of the evolutionary patterns uncovered here. Collectively, our work establishes *G. elata* as a powerful model for dissecting reductive genome evolution and reveals how obligate plant–fungus symbioses are maintained through coordinated gene loss, functional innovation, and metabolic adaptation.

## Conclusion

Our integrated pan-genome and multi-omics analyses of *Gastrodia elata* provide fundamental insights into the genomic basis of obligate plant-fungus symbiosis and reductive genome evolution. The chromosome-level genome assembly (1.09 Gb) and pan-genome construction from 12 *Gastrodia* accessions reveal extensive hybridization across cultivated germplasm, challenging conventional taxonomy based on stem color and instead supporting tuber morphology as a more reliable classification criterion among the sampled accessions. This finding has immediate implications for germplasm conservation and breeding programs of this economically important medicinal species.

Three key discoveries emerge from our work. First, the expanded *GAFP* gene family, particularly the recently evolved Class 1 lineage with modified domain architecture and elevated expression, represents a molecular innovation for balancing nutrient acquisition from *Armillaria* symbionts with defense against fungal overgrowth. Second, contrary to expectations, no horizontal gene transfer from symbiotic fungi was detected despite millions of years of intimate association, indicating that *G. elata*–fungus mutualism is maintained through metabolic exchange rather than genetic integration. Third, we delineate a comprehensive nutrient acquisition system wherein glucose from fungal trehalose hydrolysis serves as the primary carbon source, transported via an expanded STP family, while organic nitrogen is assimilated through amino acid transporters and OPT family peptide transporters coupled with urease-mediated conversion—a strategy necessitated by the loss of nitrate utilization genes.

Together, these findings establish *G. elata* as a model for understanding the genomic and metabolic adaptations underlying mycoheterotrophy, while providing essential resources for the sustainable utilization of this traditional medicinal plant.

## Material and Methods

### Plant material and public data

Experimental materials were collected from Qixingguan District, Bijie City, Guizhou Province, China. These samples were identified as four forms of *G. elata*, each displaying unique stem colors: (1) *G. elata* f. *elata* with dark red stems (TMSR) and light red stems (TMQR); (2) *G. elata* f. *viridis* with dark green stems (TMSG) and light green stems (TMQG); (3) *G. elata* f. *glauca* with dark brown stems (TMSB) and light brown stems (TMQB); and (4) *G. elata* f. *flavida* with yellow stems (TMQY) (Fig. 1A). Tender stems of the dark-red G*. elata* form (TMSR) were used for chromosomal-level whole-genome sequencing, while the remaining six forms underwent genomic resequencing. Additionally, the stems of all forms, except for yellow *G. elata* (TMQY), were subjected to transcriptome sequencing. Furthermore, transcriptome sequencing was also performed on dark-red G*. elata* (TMSR) samples at various growth and developmental stages, including immature tubers (TMMM), developing (or maturing) tubers (TMBM), mature tubers (TMJM), flowers and stems (TMHJ), and capsules (TMGS) (Fig. 1B). Each transcriptome sequencing sample included three biological replicates. We also acquired published genomic data for *G. elata* f. *glauca* from (2018) ^61^, (2020) ^4^, (2021) ^5^, as well as *G. elata* data from (2022) ^6^, from the National Center for Biotechnology Information (NCBI). Notably, the *G. elata* genome assembled by Bae et al. lacked annotation files and was newly annotated in this study^6^ (Supplementary Table 2). Coding DNA sequence (CDS) and general feature format (GFF) files corresponding to 51 green plant species were downloaded from 13 databases. Specifically, 19 CDS datasets were obtained from the Phytozome and NCBI databases (Supplementary Table 36). These 51 CDS files encompassed 45 angiosperms, two bryophyte species, two algae species, one fern species, and one gymnosperm species.

### Genome sequencing, assembly, and annotation of dark-red G*. elata*

DNA sequencing was performed using the Illumina NovaSeq 6000 at 128.41x depth, followed by paired-end sequencing on HiSeq. A 10 Gb DNA data yield with ∼15 kb SMRTbell insert-size library was prepared per PacBio protocol and sequenced on the PacBio Sequel II. Genome assembly was performed using Hifiasm (0.16.1)^13^, enhanced by Hi-C technology and ALLHiC^69^ to achieve chromosome-level resolution, with Juicebox software^70^ facilitating manual adjustments. Assembly integrity was evaluated using BUSCO (v5.4.7)^71^ and CEGMA (v2.5)^72^. Accuracy was assessed by aligning small fragment library reads to the assembly with Burrows-Wheeler Aligner (BWA) (v0.7.17-r1188)^73^, considering metrics like mapping rate, coverage, and depth distribution. Merqury (v1.4.1)^74^ was used to analyze consensus quality and completeness. Repeat sequences were annotated by combining homology alignment with *de novo* prediction using Repbase^75^ and RepeatMasker supported by LTR_FINDER, RepeatScout, and RepeatModeler.

For gene annotation, homologous protein sequences from the Ensembl and NCBI were aligned to the dark-red G*. elata* genome using TBLASTN (v2.2.26; E-value ≤ 1e-5)^76^ and GeneWise (v2.4.1)^77^ for gene structure prediction. *Ab initio* methods, including Augustus (v3.2.3)^78^, Geneid (v1.4)^79^, GENSCAN (v1.0)^80^, GlimmerHMM (v3.04)^81^ and SNAP (2013-11-29)^82^, were utilized for automated gene prediction. RNA-Seq reads from various tissues of dark-red G*. elata* were aligned to the genome with HISAT (v2.0.4)^83^ and TopHat (v2.0.11)^84^, and Stringtie (v1.3.3)^85^ and Cufflinks (v2.2.1)^86^ were used for transcript assembly. Protein function was assigned by aligning sequences with the Swiss-Prot database using BLASTP (v2.2.26; E-value ≤ 1e-5) and annotated with InterProScan (v5.31)^87^. ncRNA annotation involved tRNA prediction using tRNAscan-SE^88^ and miRNA/snRNA prediction using INFERNAL^89^ in Rfam (v14.1).

### Genome annotation of *G. elata* (2022)

To annotate the genome of *G. elata* (2022), we utilized RepeatModeler (v2.0.1)^90^ to construct a library of repeats and LTR_retriever (v2.9.0) to identify LTR retrotransposons. These repeats were then annotated in the *G. elata* (2022)^6^ genome using RepeatMasker (v4.0.9). ISO-Seq data was processed and subsequently assembled with StringTie (v1.3.3)^85^. MAKER (v3.01.03)^91^ in “est2genome” mode was employed for genome annotation, incorporating full-length transcripts from ISO-Seq and SNAP^82^ predictions, along with homology-based predictions using reference sequences from related species *Apostasia shenzhenica* (GCA_002786265.1)^26^, *Dendrobium catenatum* (GCA_001605985.2)^2^, and *Phalaenopsis equestris* (GCA_001263595.1)^92^, and two published CDS sequences of *G. elata* (*GWHAAEX00000000* and *GWHBDNU00000000*^5, 61^). Finally, Gene Model Mapper (GeMoMa)^93^ was used to integrate and analyze transcripts and predicted contents to identify protein-coding genes.

### Pangenome construction, assembly, and assessment of *Gastrodia*

The pangenome was constructed using 12 genome datasets, which include the genome data of dark-red G*. elata*, resequencing data of six *G. elata* forms from this study, as well as published genomic data of four *G. elata* and one *G. menghaiensis*. First, reads were cleaned using Trimmomatic (v0.39)^94^. *De novo* assembly was performed with SOAPdenovo2 (v2.04)^95^, complemented by transcriptomic data assembly with Trinity (v2.9.0)^96^ and CDS prediction using TransDecoder (v5.5.0)^97^. Redundant and short sequences were removed by CD-HIT (v4.8.1)^98^. Assembly optimization involved BLASTP comparison^99^ and BUSCO (v5.2.2)^100^ evaluation. OrthoFinder (v2.3.8)^101^ clustered sequences into core and pan gene families. For SNP detection, BWA^73^, SAMtools (v.1.15)^73^, and GATK (v4.3.0.0)^102^ were used for read mapping, sorting, duplicate marking, and mutation calling. Variants were filtered based on quality metrics and MAF (minor allele frequency). PCA was performed using PLINK (v1.90)^103^. INDEL sites were similarly filtered, and density plots were generated using CMplot in the R software package.

### Phylogenetic analysis of *Gastrodia* using SNP data

Initially, the SNP data was converted from VCF to PHYLIP format using the Python script “vcf2phylip”^104^. Subsequently, the phylogenetic tree was constructed using IQ-TREE (v.2.0.3)^105^. The optimal model was determined using the “ModelFinder” function based on the Bayesian Information Criterion (BIC)^106^. Finally, the maximum likelihood tree was constructed using this model, with 1,000 ultrafast bootstrap tests and 1,000 SH-aLRT (Shimodaira-Hasegawa approximate Likelihood Ratio Test) branch tests performed to ensure the robustness of the inferred phylogeny.

### Detection of hybridization signals in 11 *G. elata* samples

To detect hybridization signals in eleven samples of *G. elata*, SNP data in PHYLIP format were analyzed using the HyDe software^107^. Subsequently, a hybridization network map was generated using the R packages tidygraph, ggraph, and ggplot2.

### Genome collinearity analysis and visualization

Genomic collinearity was analyzed using MCScanX^108^ based on the protein sequences from GFF3 files of dark-red G*. elata*, *G. elata* f. *glauca* (2018), *G. elata* f. *glauca* (2021), *G. elata* (2022), and *G. menghaiensis*^109, 110^. Genome syntenic visualization was conducted using jcvi (https://github.com/tanghaibao/jcvi). Dot plots for syntenic gene pairs were generated by WGDI. Ks values of syntenic gene pairs were quantified using the Nei-Gojobori method in PAML (v.4.9h)^111^. Median and peak Ks values were determined and visually represented via WGDI.

### Phylogeny reconstruction of 54 land plants

A phylogenetic analysis of 54 land plant species was conducted using a standardized pipeline^112^. Protein sequences were subjected to BLASTP with an E-value threshold of 1e-5, followed by clustering with MCL software^113^. Amino acid sequence alignments were generated by MAFFT (v7.508)^114^ and converted to nucleic acid alignments using PAL2NAL^115^. Low-quality regions were trimmed with trimAl^116^. The final maximum likelihood tree was reconstructed using IQ-TREE (v.2.0.3)^105^ with the parameter “-m GTR+G −bb 1000”.

### Divergence time estimation of land plants

To estimate the divergence times of the 54 land plant genomes, we employed seven fossil calibrations spanning from 23.2 million years ago (Mya) to 515.5 Mya^117, 118, 119, 120, 121^ and one secondary calibration at 669.9-972.4 Mya^120^. Using 437 single-copy genes with >90% coverage and >1000 bp length, divergence times were analyzed with MCMCTREE in PAML^111^. The analysis utilized an autocorrelation model, the GTR substitution model, and a unified prior for node times. Markov chain Monte Carlo (MCMC) sampling was conducted for 500,000 steps after a burn-in of 200,000 steps, with sampling every 20 steps.

### Detection of gene family contraction, expansion and duplications

We analyzed gene family contraction and expansion using protein sequences from 54 land plant genomes. DIAMOND (v2.0.15.153)^122^ was employed for all-against-all BLASTP searches with specific parameters to obtain bit scores. These scores were then used by PhyloMCL^123^ for gene family clustering. Finally, the “dollo parsimony” module in Tree2gd software determined gene family contraction and expansion.

### Analysis of genes under positive selection in 13 orchid genomes

The protein sequences from 13 Orchidaceae genomes were clustered using OrthoFinder^101^ with default settings. MAFFT^114^ was then used to align these sequences, followed by deriving nucleotide alignments from the protein alignments with PAL2NAL^115^. Gene family trees were inferred using IQ-TREE^124^ based on these alignments. The “Ortho_Retriever” module in PhyloTracer disassembled these trees into 1,837 single-copy gene family trees. Finally, ete3 evol module from ETE 3 toolkit ^125^ analyzed these trees along with their corresponding aligned CDS sequences.

### RNA-seq data analysis

The RNA-seq analysis involved cDNA library preparation, paired-end sequencing on Illumina NovaSeq 6000, FastQC^126^ for quality control, HISAT2 (v2.0.5)^127^ for alignment to the reference genome (dark-red G*. elata*), StringTie (v1.3.3)^85^ for transcript assembly, Subread software^128^ for read counts calculation, DESeq2 (v1.20.0)^129^ for identifying DEGs (|log_2_(Fold change)| >= 1 & padj <= 0.05) ClusterProfiler for functional enrichment, and WGCNA for gene co-expression analysis.

### Molecular evolution of key immune responsive gene

The antifungal protein sequence (https://www.ncbi.nlm.nih.gov/protein/AAK59994) from *G. elata* was retrieved from NCBI and used as a BLASTP query (E-value < 1e-5) across 54 land plants. Homologous sequences were aligned using MAFFT^114^ with parameters “-maxiterate 1000 −localpair” for multiple sequence alignment. The phylogenetic tree was constructed using IQ-TREE^105^ with GTR+G model. Conserved protein structures were validated through NCBI Conserved Domains. The resulting phylogenetic tree of the antifungal protein from *G. elata* was refined using FigTree (v1.4.4) (http://tree.bio.ed.ac.uk/software/figtree/) and visualized with iTOL^130^.

### Biotin-GAFP Pull-down Experiment and LC-MS/MS Detection

The GAFP protein was extracted from fresh tubers of *G. elata* and purified using ion exchange chromatography ^12^. The collected fractions were analyzed by sodium dodecyl sulfate-polyacrylamide gel electrophoresis (SDS-PAGE) to confirm the molecular weight and purity of the protein. The total protein from *A. gallica* mycelium was extracted using a fungal mycelium protein extraction kit.

Biotinylation was performed using a biotinylation kit to covalently attach a biotin tag (Biotin-GAFP) to the purified GAFP protein. Based on the high specificity and strong affinity of the biotin-streptavidin interaction, streptavidin magnetic beads were used for pull-down experiments to capture the target protein. Two experimental groups were set up, each with three replicates. The first group included Bio-GAFP, *A. gallica* total protein, and streptavidin magnetic beads, while the second group was the blank control, containing only *A. gallica* total protein and streptavidin magnetic beads^4, 29^. The obtained target protein was dissociated into peptides, desalted, and analyzed by LC-MS/MS to obtain peptide data ^30, 31^. The data were then processed using MaxQuant (v2.6.7) ^131^ software to search the protein database (downloaded from the Uniprot website for *A. gallica* proteins). The peptide identification results were used for subsequent bioinformatics analysis.

### Molecular evolution of key enzyme genes involved in photosynthesis pathway across Viridiplantae

To investigate the molecular evolution of key enzyme genes in the photosynthesis pathway across Viridiplantae, we focused on the Rubisco activase genes and *PPC* gene family. For Rubisco activase, we used *A. thaliana* sequences from NCBI to perform homologous comparisons in *A. thaliana* (C3), *Z. mays* (C4), *K. fedtschenkoi* (CAM), and dark-red G*. elata*. Conserved domains were verified using NCBI CDD and PfamScan^132^, excluding genes lacking Rubisco domains. For *PPC* genes, we obtained the HMM profile (PF00311) from PFAM^133^ and searched across 54 land plant genomes using HMMER 3.1. PPC domain verification was conducted using NCBI CDD, excluding sequences lacking this domain. Multiple *PPC* protein sequences were aligned using MAFFT^114^, and a phylogenetic tree was constructed with IQ-TREE^105^ using the SYM + R8 model, visualized in iTOL^130^.

### Molecular evolution of antifungal protein and MADS-box gene family

To study the molecular evolution of key immune responsive genes, the antifungal protein sequence from *G. elata* was retrieved and used for BLASTP across 54 land plants. Homologous sequences were aligned using MAFFT^114^, and a phylogenetic tree was constructed using IQ-TREE^105^ with GTR+G model. The tree was refined using FigTree and visualized with iTOL^130^. For the MADS-box gene family, HMM file obtained from PFAM^133^ was used with HMMER 3.1 to identify homologs in *G. elata* and *A. thaliana* genomes, which were then aligned using MAFFT^114^. Phylogenetic analysis was conducted using IQ-TREE^105^ with the SYM+R8 model, and the tree was visualized using FigTree and iTOL^130^.

### Identification and evolutionary characterization of HGT-acquired genes in Orchidaceae plants

This study employed a systematic multi-step pipeline to identify potential HGT-acquired genes in Orchidaceae plants, comprising three key stages: Initial screening using HGTphyloDetect^134^ and AvP^135^; Multi-dimensional filtering; and Phylogenetic validation.

First, DIAMOND BLASTp (-evalue 1e-5; -max_target_seqs 5,000,000) was used to align protein sequences from 13 orchid genomes against the Non-Redundant Protein Database (NR, https://ftp.ncbi.nlm.nih.gov/blast/db/FASTA/nr.gz), the OneKP database^136^ and the 13 orchid genomes themselves. The resulting alignments served as inputs for HGT detection tools. Candidate HGT-acquired genes were initially identified using both AvP and HGTphyloDetect. To ensure reliability, strict filtering criteria were applied: (1) Removal of genes homologous to rice and *Arabidopsis thaliana* endogenous viruses (e-value ≤ 1e-5). (2) Removal of sequences similar to rice and *A. thaliana* mitochondrial/chloroplast genomes (e-value ≤ 1e-5). (3) BLASTp matches for remaining candidates were categorized into seven groups (Bacteria, Fungi, Archaea, Viruses, Metazoa, Viridiplantae, Other_Eukaryota). Groups with ≤ 2 species were excluded as likely contaminants. Each candidate was required to match ≥ 2 orchid species to eliminate sequencing artifacts. Phylogenetic validation: for each candidate HGT-acquired gene, homologous sequences were retrieved and curated: For genes with ≤100 homologous sequences, the top BLAST hit per species was selected; For genes with >100 homologous sequences, the top hit per family was selected. Multiple sequence alignment was performed using MAFFT (--auto)^114^, followed by maximum-likelihood tree construction with FastTree (-lg -gamma)^137^. Reliable HGT events were defined by: Paraphyletic donor lineages and monophyletic plant clades.

RNA-seq Mapping: high-quality genomes of five Armillaria and five Mycena species were downloaded from the Global Catalogue of Fungi Genomes (https://globalfungi.com) as reference genomes (Supplementary Fig.13). BBsplit^138^ was used to map 17 RNA-seq datasets to these fungal genomes. To explore functional implications, PPI (Protein-protein interaction) networks of HGT-acquired genes were analyzed: *Arabidopsis thaliana* PPI data were downloaded from STRING v12.0 (https://stringdb-downloads.org/download/protein.physical.links.full.v12.0/3702.protein.physical.links.full.v12.0.txt.gz). Transcription factor (TF) genes were obtained from TAIR (https://www.arabidopsis.org/ ), and their PPI networks (confidence score ≥ 700) were extracted. PPI networks were visualized using Cytoscape v3.10.3^139^, and node degrees were calculated via the “Network Analyzer” module. For HGT-acquired genes lacking direct STRING coverage, homologous networks were inferred using the STRING “Protein by Sequence” tool and their PPI networks (confidence score ≥ 700) were extracted. Functional enrichment analysis was performed with the ClueGO plugin (v2.5.10) in Cytoscape^140^.

### Molecular evolution of transporter protein genes in *G. elata*

To investigate the molecular evolution mechanism of nutrient-related genes in *G. elata*, this study downloaded the CDS sequences of monosaccharide transporters, sugar metabolism-related genes, amino acid transporters, nitrogen-related genes, fatty acid metabolism-related genes, and transporter proteins from the TAIR database^51, 54, 55, 59, 65, 141^ (Supplementary Table 36). In addition, using the sequences of *AsUGT*^36^ as the query genes for key enzymes in gastrodin synthesis^112^ (Supplementary Table 36). BLASTP alignment was performed against the genomes of 54 terrestrial plant species (with e-value: 1e-5). Gene family clustering was conducted using MCL^113^, and multiple sequence alignment was performed using MAFFT^114^, with the alignment results converted into nucleotide alignments. Phylogenetic trees were constructed using IQ-TREE^105^ (under the SYM + R8 model) and the root positions of the trees were determined using Tree2gd^142^. Furthermore, the Python toolkit ete3 was used to split the superfamily gene trees of monosaccharide transporters and amino acid transporters, ensuring each gene ID belonged to only one subfamily. Finally, the phylogenetic trees were visualized using the Tree Visualizer tool (https://github.com/YiyongZhao/PhyloTracer).

In this study, some homologous genes in *G. elata* were selected, and their expression levels in *G. elata* at the immature tuber, developing tuber, mature tuber, flower and stem, and capsule stages were presented via heatmaps. The data were first used to calculate the average value of each stage, and then transformed using log₂(FPKM+1) to eliminate heteroscedasticity; the original data of other homologous genes are available in the Supplementary Table. Meanwhile, *G. elata* samples were divided into three groups: the first group included the tuber stages (immature tuber, developing tuber, mature tuber); the second group included flower and stem; the third group included the fruit stage. Based on these groupings, cluster analysis was used to generate plots for displaying the differences in gene expression levels among groups^143^. In addition, the copy numbers of the aforementioned genes in *Gastrodia* species, Orchidaceae plants, Monocots, and eudicots were presented in the Supplementary Table.

## Data availability statement

The sequencing data utilized in this study have been deposited at the National Center for Biotechnology Information (NCBI) under project number PRJNA941105 (https://dataview.ncbi.nlm.nih.gov/object/PRJNA941105?reviewer=v7gtpai9dbbnqedajb0n5vk5a). This dataset comprises next-generation sequencing data, third-generation sequencing data, Hi-C data, whole genome sequencing data of the dark-red G*. elata* genome, transcriptomic data from various stages of the dark-red G*. elata*, as well as transcriptomic data and whole genome resequencing data of *G. elata* in different colors. Requests for additional datasets and scripts should be directed to the corresponding author, Mingjin Huang. The phylogenetic tree constructed based on candidate HGT-acquired genes has been uploaded to FigShare (https://doi.org/10.6084/m9.figshare.16831776).

## Contribution

Mingjin Huang and Yiyong Zhao designed the study. Mingjin Huang, Cheng Li, and Dachang Wang prepared the plant samples and performed sequencing. Shanshan Luo, Linshuang Tang, Hao Yin, Yiyong Zhao, Mingjin Huang, Daliang Liu, Mengge Li, Zhipeng Li, Qiyu Chen, Yinjie Jiao, Tao Li, and Yanlin Hao conducted the experiments and analyzed the data. Mingjin Huang, Yiyong Zhao, Shanshan Luo, Linshuang Tang and Hao Yin drafted the manuscript. Yiyong Zhao, Mingjin Huang, Lin Cheng, Cheng Li, Hualei Wang, Guojin Zhang, Wei Wang, Ruimin Wang, Dandan Li, Jin He and Xinhao Sun revised the manuscript. All authors approved the final version of the manuscript.

## Acknowledgements

We thank the anonymous reviewers for their valuable comments on our manuscript. We are deeply appreciative of the State Key Laboratory of Big Data at Guizhou University for providing supercomputing services essential for data analysis. This research was supported by funding from the Traditional Chinese Medicine Industry Development Project of Guizhou Province, China (Nos. [2019]175, [2020]101, [2021]145); Demonstration of Breeding, Propagation, and High-Yield & Stable Production Technology Enhancement for New Varieties of *G. elata* (Herbal Medicine Cluster Project of the Guizhou Provincial Department of Agriculture and Rural Affairs, 2025); Guizhou Chuangheyuan Technology Co., Ltd (No. [2021]001); the Construction Project of Modern Industry Technology System for Traditional Chinese Medicinal Materials in Guizhou Province, China (No. GZCYTX-02, 2024); and the Key Laboratory of High Quality, High Efficiency, and Yield Enhancement in Grain and Oil Crops (Qian-Ke-He-Platform ZSYS[2025] 037).

## Supplementary figure legends

**Figure S1.** Overview of genome assembly, analysis, and sequencing depth

A. Workflow of genome assembly.

B. Base composition statistics of dark-red G*. elata*.

C. BUSCO assessment of genome completeness.

D. Sequencing depth distribution.

**Figure S2.** Transposable element (TE) sequence divergence and gene function annotation in the dark-red G*. elata* genome.

A. TE sequence divergence distribution: The x-axis shows divergence degrees between TEs annotated in the dark-red G*. elata* genome and Repbase reference sequences. The y-axis represents the proportion of TEs with specific divergence degrees. Colors distinguish different TE types.

B. Gene function annotation summary: By aligning known protein databases with predicted protein sequences from the dark-red G*. elata* genome, 90.57% of genes were functionally annotated.

**Figure S3.** Quantitative genetic analysis of different color forms of *G. elata*

A. RNA-seq gene expression quantification, adjusted for sequencing depth and gene length using FPKM (Fragments Per Kilobase Million), illustrated by box plots showing expression levels across samples. X-axis: sample names; Y-axis: log_2_(FPKM + 1).

B. Heatmap depicting correlation coefficients between samples based on FPKM values, highlighting group differences and sample repetition.

C. PCA plot evaluating inter-group differences and intra-group consistency using gene expression values (FPKM). Samples between groups ideally disperse, while within-group samples cluster.

D. Co-expression Venn diagram showing uniquely expressed genes per sample and co-expressed genes across samples.

TMSG: dark green *G. elata*, TMQG: light green *G. elata*, TMSR: dark-red G*. elata*, TMQR: light red *G. elata*, TMSB: dark brown *G. elata*, TMQB: light brown *G. elata*.

**Figure S4.** Histogram of the number of differentially expressed genes of different color stems for *G. elata*.

TMSG: dark green *G. elata*, TMQG: light green *G. elata*, TMSR: dark-red G *elata*, TMQR: light red *G. elata*, TMSB: dark brown *G. elata*, TMQB: light brown *G. elata*.

**Figure S5.** GO and KEGG functional enrichment analyses in different color stems of *G. elata* transcriptome

A. Scatter plots displaying the top 30 statistically significant GO terms from GO enrichment analysis.

B. Scatter plots showcasing the 20 most significant KEGG pathways from KEGG enrichment analysis. The horizontal axis represents sample comparisons, and the vertical axis lists GO terms/pathways. Point size reflects the number of annotated genes, and the color gradient (from red to purple) indicates enrichment significance.

**Figure S6.** Cluster analysis of 13 volatile compounds detected by gas chromatography-mass spectrometry (GC-MS) in various germplasms of *Gastrodia elata*. The x-axis is the *Gastrodia elata* sample, and the y-axis is the name of the volatile compound. The label on the right is the correlation between the compound and *Gastrodia elata*, blue represents low correlation, and red represents high correlation.

**Figure S7.** Box plots (A, B, C) showing protein sequence similarity comparisons among 11 *G. elata* forms and one *G. menghaiensis*. Labels A to L represent: dark green *G. elata*, dark brown *G. elata*, *G. elata* f. *glauca* (2020), *G. elata* f. *glauca* (2021), *G. elata* f. *glauca* (2018), light brown *G. elata*, *G. elata* (2022), light green *G. elata*, light red *G. elata*, *G. elata* f. *flavida*, dark-red G*. elata* (this study), and *G. menghaiensis*, respectively. Each comparison (e.g., C/L) indicates a BLASTP analysis of protein sequences between varieties C and L, with bitscores tallied, de-weighted, and normalized using log10 transformation. A total of 66 comparisons are shown, sorted by the median value of similarity scores, to evaluate protein sequence similarity across the 12 varieties.

**Figure S8.** Ks dot plot analysis of four *G. elata* forms (A: *G. elata f. glauca* 2021, B: *G. elata* 2022, C: *G. menghaiensis*, D: dark-red G*. elata*). Colors represent Ks values of homologous gene pairs in covariant genomic segments, illustrating intragenomic covariance.

**Figure S9.** The phylogenetic tree comprises *Gastrodia* and multiple species. Dicotyledonous plants are denoted in green font, *Gastrodia*’s position is shown in blue font alongside its corresponding group and class 1 antifungal proteins are highlighted in red font.

**Figure S10.** A phylogenetic tree of *AsUGT* genes from 54 land plant species. The involved plants include 54 plants such as *Gastrodia* and *Dendrobium* (Supplementary Table 20). The vertical columns from left to right in the figure represent gene ID, species name, species family, and expression level, respectively. Interspecific and intraspecific whole-genome duplication events are marked in the upper left corner of the figure. Bootstrap values have been labeled on the phylogenetic tree, which indicate the credibility of the branches. The query genes used in *A. thaliana* have been marked on the tree. The right side of the figure shows the expression data of duplicated genes in dark-red G*. elata*, covering the following developmental stages: TMMM (immature tuber), TMBM (developing tuber), TMJM (mature tuber), TMGS (capsule), and TMHJ (inflorescence). Each stage has three biological replicates, and the data have been averaged.

**Figure S11.** Phylogenetic tree of nine species, including dark-red G*. elata*. Colors: C3/CAM (yellow), C4 (blue), CAM (red).

**Figure S12.** Expression of the *Gae* gene across different color forms of *G. elata* and various developmental stages of TMSR.

A. Heat map illustrating the expression pattern of the *Gae* gene in six distinct color forms of *Gastrodia elata*, with the bacterial *PPC* gene (BTPC) indicated in blue. The key gene, *K. fedtschenkoi*, is denoted by a red pentagram. “*K. fedtschenkoi* Leaf_1” represents mature leaves of 1. K. fedtschen-koi, while “K. fedtschenkoi Leaf_2” signifies 4-week-old young leaves. “*K. fedtschenkoi* Root_1” refers to the root of *K. fedtschenkoi*. Similarly, “*A. thaliana* Leaf_3” and “*A. thaliana* Root_2” denote mature leaves and roots of *A. thaliana*, respectively. TMQR: light red *G. elata*, TMSR: dark-red G*. elata*, TMQG: light green *G. elata*, TMSG: dark green *G. elata*, TMQB: light brown *G. elata*, TMSB: dark brown *G. elata*.

B. Heat map consisting of the expression of the *Gae* gene during the five stages of dark-red G*. elata*. TMMM: immature tuber, TMBM: developing tuber; TMJM: mature tuber, TMHJ: inflorescence, TMGS: capsule.

**Figure S13.** Chlorophyll biosynthesis pathway in plants and expression of related enzyme genes in *G. elata*. Heatmap (right) shows expression levels of chlorophyll biosynthesis-related enzyme genes. *HEMA*: Glutamyl-tRNA reductase*. GSA*: Glutamate-1-semialdehyde 2, 1-aminomutase*. HEMB*: 5-aminolevulinate dehydratase*. HEMC*: Porphobilinogen deaminase*. HEMD*: Uroporphyrinogen III synthase*. UROS*: Uroporphyrinogen III decarboxylase*. HEMF*: Coproporphyrinogen III oxidase*. HEMG*: Protoporphyrinogen IX oxidase. *CHLD*: Magnesium chelatase subunit D*. CHLM*: Magnesium-protoporphyrin IX methyltransferase*. DVR*: Divinyl reductase*. POR*: Protochlorophyllide oxidoreductase*. CHLG*: Chlorophyll synthase*. CAO*: Chlorophyllide a oxygenase.

**Figure S14.** Workflow for HGT identification. The HGT identification process in this study includes three main steps: (1) Preliminary screening of candidate HGT genes; (2) Multi-dimensional filtering; (3) Phylogenetic validation. Yellow oval boxes represent protein sequences, green boxes indicate BLAST result files, blue oval boxes denote the candidate HGT-acquired gene sets obtained at each filtering step, and red boxes represent the judgment criteria. The text on the arrows between each analytical step indicates the corresponding screening process.

**Figure S15.** Degree distributions in the protein-protein interaction network between HGT-acquired genes and transcription factor genes.

**Figure S16.** A. It shows the domain map of 26 *A. gallica* proteins, with different colors representing different domains, and the horizontal axis indicating the length of the proteins. B. It presents the GO enrichment analysis of the 26 *A. gallica* proteins. C. It displays the KEGG enrichment analysis of the 26 *A. gallica* proteins.

**Figure S17.** A phylogenetic tree of *NST* genes from 54 land plant species. The involved plants include 54 plants such as *Gastrodia* and *Dendrobium* (Supplementary Table 20). The vertical columns from left to right in the figure represent gene ID, species name, species family, and expression level, respectively. Interspecific and intraspecific whole-genome duplication events are marked in the upper left corner of the figure. Bootstrap values have been labeled on the phylogenetic tree, which indicate the credibility of the branches. The query genes used in *A. thaliana* have been marked on the tree. The right side of the figure shows the expression data of duplicated genes in dark-red G*. elata*, covering the following developmental stages: TMMM (immature tuber), TMBM (developing tuber), TMJM (mature tuber), TMGS (capsule), and TMHJ (inflorescence). Each stage has three biological replicates, and the data have been averaged.

**Figure S18.** A phylogenetic tree of *INT* genes from 54 land plant species. The involved plants include 54 plants such as *Gastrodia* and *Dendrobium* (Supplementary Table 20). The vertical columns from left to right in the figure represent gene ID, species name, species family, and expression level, respectively. Interspecific and intraspecific whole-genome duplication events are marked in the upper left corner of the figure. Bootstrap values have been labeled on the phylogenetic tree, which indicate the credibility of the branches. The query genes used in *A. thaliana* have been marked on the tree. The right side of the figure shows the expression data of duplicated genes in dark-red G*. elata*, covering the following developmental stages: TMMM (immature tuber), TMBM (developing tuber), TMJM (mature tuber), TMGS (capsule), and TMHJ (inflorescence). Each stage has three biological replicates, and the data have been averaged.

**Figure S19.** A phylogenetic tree of *STP* genes from 54 land plant species. The involved plants include 54 plants such as *Gastrodia* and *Dendrobium* (Supplementary Table 20). The vertical columns from left to right in the figure represent gene ID, species name, species family, and expression level, respectively. Interspecific and intraspecific whole-genome duplication events are marked in the upper left corner of the figure. Bootstrap values have been labeled on the phylogenetic tree, which indicate the credibility of the branches. The query genes used in *A. thaliana* have been marked on the tree. The right side of the figure shows the expression data of duplicated genes in dark-red G*. elata*, covering the following developmental stages: TMMM (immature tuber), TMBM (developing tuber), TMJM (mature tuber), TMGS (capsule), and TMHJ (inflorescence). Each stage has three biological replicates, and the data have been averaged.

**Figure S20.** A phylogenetic tree of *PGLCT* genes from 54 land plant species. The involved plants include 54 plants such as *Gastrodia* and *Dendrobium* (Supplementary Table 20). The vertical columns from left to right in the figure represent gene ID, species name, species family, and expression level, respectively. Interspecific and intraspecific whole-genome duplication events are marked in the upper left corner of the figure. Bootstrap values have been labeled on the phylogenetic tree, which indicate the credibility of the branches. The query genes used in *A. thaliana* have been marked on the tree. The right side of the figure shows the expression data of duplicated genes in dark-red G*. elata*, covering the following developmental stages: TMMM (immature tuber), TMBM (developing tuber), TMJM (mature tuber), TMGS (capsule), and TMHJ (inflorescence). Each stage has three biological replicates, and the data have been averaged.

**Figure S21.** A phylogenetic tree of *VGT* genes from 54 land plant species. The involved plants include 54 plants such as *Gastrodia* and *Dendrobium* (Supplementary Table 20). The vertical columns from left to right in the figure represent gene ID, species name, species family, and expression level, respectively. Interspecific and intraspecific whole-genome duplication events are marked in the upper left corner of the figure. Bootstrap values have been labeled on the phylogenetic tree, which indicate the credibility of the branches. The query genes used in *A. thaliana* have been marked on the tree. The right side of the figure shows the expression data of duplicated genes in dark-red G*. elata*, covering the following developmental stages: TMMM (immature tuber), TMBM (developing tuber), TMJM (mature tuber), TMGS (capsule), and TMHJ (inflorescence). Each stage has three biological replicates, and the data have been averaged.

**Figure S22.** A phylogenetic tree of *TMT* genes from 54 land plant species. The involved plants include 54 plants such as *Gastrodia* and *Dendrobium* (Supplementary Table 20). The vertical columns from left to right in the figure represent gene ID, species name, species family, and expression level, respectively. Interspecific and intraspecific whole-genome duplication events are marked in the upper left corner of the figure. Bootstrap values have been labeled on the phylogenetic tree, which indicate the credibility of the branches. The query genes used in *A. thaliana* have been marked on the tree. The right side of the figure shows the expression data of duplicated genes in dark-red G*. elata*, covering the following developmental stages: TMMM (immature tuber), TMBM (developing tuber), TMJM (mature tuber), TMGS (capsule), and TMHJ (inflorescence). Each stage has three biological replicates, and the data have been averaged.

**Figure S23.** A phylogenetic tree of *trehalase* genes from 54 land plant species. The involved plants include 54 plants such as *Gastrodia* and *Dendrobium* (Supplementary Table 20). The vertical columns from left to right in the figure represent gene ID, species name, species family, and expression level, respectively. Interspecific and intraspecific whole-genome duplication events are marked in the upper left corner of the figure. Bootstrap values have been labeled on the phylogenetic tree, which indicate the credibility of the branches. The query genes used in *A. thaliana* have been marked on the tree. The right side of the figure shows the expression data of duplicated genes in dark-red G*. elata*, covering the following developmental stages: TMMM (immature tuber), TMBM (developing tuber), TMJM (mature tuber), TMGS (capsule), and TMHJ (inflorescence). Each stage has three biological replicates, and the data have been averaged.

**Figure S24.** A phylogenetic tree of *sweet* genes from 54 land plant species. The involved plants include 54 plants such as *Gastrodia* and *Dendrobium* (Supplementary Table 20). The vertical columns from left to right in the figure represent gene ID, species name, species family, and expression level, respectively. Interspecific and intraspecific whole-genome duplication events are marked in the upper left corner of the figure. Bootstrap values have been labeled on the phylogenetic tree, which indicate the credibility of the branches. The query genes used in *A. thaliana* have been marked on the tree. The right side of the figure shows the expression data of duplicated genes in dark-red G*. elata*, covering the following developmental stages: TMMM (immature tuber), TMBM (developing tuber), TMJM (mature tuber), TMGS (capsule), and TMHJ (inflorescence). Each stage has three biological replicates, and the data have been averaged.

**Figure S25.** A phylogenetic tree of *SUS* genes from 54 land plant species. The involved plants include 54 plants such as *Gastrodia* and *Dendrobium* (Supplementary Table 20). The vertical columns from left to right in the figure represent gene ID, species name, species family, and expression level, respectively. Interspecific and intraspecific whole-genome duplication events are marked in the upper left corner of the figure. Bootstrap values have been labeled on the phylogenetic tree, which indicate the credibility of the branches. The query genes used in *A. thaliana* have been marked on the tree. The right side of the figure shows the expression data of duplicated genes in dark-red G*. elata*, covering the following developmental stages: TMMM (immature tuber), TMBM (developing tuber), TMJM (mature tuber), TMGS (capsule), and TMHJ (inflorescence). Each stage has three biological replicates, and the data have been averaged.

**Figure S26.** A phylogenetic tree of *Inv* genes from 54 land plant species. The involved plants include 54 plants such as *Gastrodia* and *Dendrobium* (Supplementary Table 20). The vertical columns from left to right in the figure represent gene ID, species name, species family, and expression level, respectively. Interspecific and intraspecific whole-genome duplication events are marked in the upper left corner of the figure. Bootstrap values have been labeled on the phylogenetic tree, which indicate the credibility of the branches. The query genes used in *A. thaliana* have been marked on the tree. The right side of the figure shows the expression data of duplicated genes in dark-red G*. elata*, covering the following developmental stages: TMMM (immature tuber), TMBM (developing tuber), TMJM (mature tuber), TMGS (capsule), and TMHJ (inflorescence). Each stage has three biological replicates, and the data have been averaged.

**Figure S27.** A phylogenetic tree of *SUT* genes from 54 land plant species. The involved plants include 54 plants such as *Gastrodia* and *Dendrobium* (Supplementary Table 20). The vertical columns from left to right in the figure represent gene ID, species name, species family, and expression level, respectively. Interspecific and intraspecific whole-genome duplication events are marked in the upper left corner of the figure. Bootstrap values have been labeled on the phylogenetic tree, which indicate the credibility of the branches. The query genes used in *A. thaliana* have been marked on the tree. The right side of the figure shows the expression data of duplicated genes in dark-red G*. elata*, covering the following developmental stages: TMMM (immature tuber), TMBM (developing tuber), TMJM (mature tuber), TMGS (capsule), and TMHJ (inflorescence). Each stage has three biological replicates, and the data have been averaged.

**Figure S28.** A phylogenetic tree of *AMY* genes from 54 land plant species. The involved plants include 54 plants such as *Gastrodia* and *Dendrobium* (Supplementary Table 20). The vertical columns from left to right in the figure represent gene ID, species name, species family, and expression level, respectively. Interspecific and intraspecific whole-genome duplication events are marked in the upper left corner of the figure. Bootstrap values have been labeled on the phylogenetic tree, which indicate the credibility of the branches. The query genes used in *A. thaliana* have been marked on the tree. The right side of the figure shows the expression data of duplicated genes in dark-red G*. elata*, covering the following developmental stages: TMMM (immature tuber), TMBM (developing tuber), TMJM (mature tuber), TMGS (capsule), and TMHJ (inflorescence). Each stage has three biological replicates, and the data have been averaged.

**Figure S29.** A phylogenetic tree of *MGAM* genes from 54 land plant species. The involved plants include 54 plants such as *Gastrodia* and *Dendrobium* (Supplementary Table 20). The vertical columns from left to right in the figure represent gene ID, species name, species family, and expression level, respectively. Interspecific and intraspecific whole-genome duplication events are marked in the upper left corner of the figure. Bootstrap values have been labeled on the phylogenetic tree, which indicate the credibility of the branches. The query genes used in *A. thaliana* have been marked on the tree. The right side of the figure shows the expression data of duplicated genes in dark-red G*. elata*, covering the following developmental stages: TMMM (immature tuber), TMBM (developing tuber), TMJM (mature tuber), TMGS (capsule), and TMHJ (inflorescence). Each stage has three biological replicates, and the data have been averaged.

**Figure S30.** A phylogenetic tree of *MEX* genes from 54 land plant species. The involved plants include 54 plants such as *Gastrodia* and *Dendrobium* (Supplementary Table 20). The vertical columns from left to right in the figure represent gene ID, species name, species family, and expression level, respectively. Interspecific and intraspecific whole-genome duplication events are marked in the upper left corner of the figure. Bootstrap values have been labeled on the phylogenetic tree, which indicate the credibility of the branches. The query genes used in *A. thaliana* have been marked on the tree. The right side of the figure shows the expression data of duplicated genes in dark-red G*. elata*, covering the following developmental stages: TMMM (immature tuber), TMBM (developing tuber), TMJM (mature tuber), TMGS (capsule), and TMHJ (inflorescence). Each stage has three biological replicates, and the data have been averaged.

**Figure S31.** A phylogenetic tree of *Chi* genes from 54 land plant species. The involved plants include 54 plants such as *Gastrodia* and *Dendrobium* (Supplementary Table 20). The vertical columns from left to right in the figure represent gene ID, species name, species family, and expression level, respectively. Interspecific and intraspecific whole-genome duplication events are marked in the upper left corner of the figure. Bootstrap values have been labeled on the phylogenetic tree, which indicate the credibility of the branches. The query genes used in *A. thaliana* have been marked on the tree. The right side of the figure shows the expression data of duplicated genes in dark-red G*. elata*, covering the following developmental stages: TMMM (immature tuber), TMBM (developing tuber), TMJM (mature tuber), TMGS (capsule), and TMHJ (inflorescence). Each stage has three biological replicates, and the data have been averaged.

**Figure S32.** A phylogenetic tree of *AGAL* genes from 54 land plant species. The involved plants include 54 plants such as *Gastrodia* and *Dendrobium* (Supplementary Table 20). The vertical columns from left to right in the figure represent gene ID, species name, species family, and expression level, respectively. Interspecific and intraspecific whole-genome duplication events are marked in the upper left corner of the figure. Bootstrap values have been labeled on the phylogenetic tree, which indicate the credibility of the branches. The query genes used in *A. thaliana* have been marked on the tree. The right side of the figure shows the expression data of duplicated genes in dark-red G*. elata*, covering the following developmental stages: TMMM (immature tuber), TMBM (developing tuber), TMJM (mature tuber), TMGS (capsule), and TMHJ (inflorescence). Each stage has three biological replicates, and the data have been averaged.

**Figure S33.** A phylogenetic tree of *AAP* genes from 54 land plant species. The involved plants include 54 plants such as *Gastrodia* and *Dendrobium* (Supplementary Table 20). The vertical columns from left to right in the figure represent gene ID, species name, species family, and expression level, respectively. Interspecific and intraspecific whole-genome duplication events are marked in the upper left corner of the figure. Bootstrap values have been labeled on the phylogenetic tree, which indicate the credibility of the branches. The query genes used in *A. thaliana* have been marked on the tree. The right side of the figure shows the expression data of duplicated genes in dark-red G*. elata*, covering the following developmental stages: TMMM (immature tuber), TMBM (developing tuber), TMJM (mature tuber), TMGS (capsule), and TMHJ (inflorescence). Each stage has three biological replicates, and the data have been averaged.

**Figure S34.** A phylogenetic tree of *LHT* genes from 54 land plant species. The involved plants include 54 plants such as *Gastrodia* and *Dendrobium* (Supplementary Table 20). The vertical columns from left to right in the figure represent gene ID, species name, species family, and expression level, respectively. Interspecific and intraspecific whole-genome duplication events are marked in the upper left corner of the figure. Bootstrap values have been labeled on the phylogenetic tree, which indicate the credibility of the branches. The query genes used in *A. thaliana* have been marked on the tree. The right side of the figure shows the expression data of duplicated genes in dark-red G*. elata*, covering the following developmental stages: TMMM (immature tuber), TMBM (developing tuber), TMJM (mature tuber), TMGS (capsule), and TMHJ (inflorescence). Each stage has three biological replicates, and the data have been averaged.

**Figure S35.** A phylogenetic tree of *CAT* genes from 54 land plant species. The involved plants include 54 plants such as *Gastrodia* and *Dendrobium* (Supplementary Table 20). The vertical columns from left to right in the figure represent gene ID, species name, species family, and expression level, respectively. Interspecific and intraspecific whole-genome duplication events are marked in the upper left corner of the figure. Bootstrap values have been labeled on the phylogenetic tree, which indicate the credibility of the branches. The query genes used in *A. thaliana* have been marked on the tree. The right side of the figure shows the expression data of duplicated genes in dark-red G*. elata*, covering the following developmental stages: TMMM (immature tuber), TMBM (developing tuber), TMJM (mature tuber), TMGS (capsule), and TMHJ (inflorescence). Each stage has three biological replicates, and the data have been averaged.

**Figure S36.** A phylogenetic tree of *GAT* genes from 54 land plant species. The involved plants include 54 plants such as *Gastrodia* and *Dendrobium* (Supplementary Table 20). The vertical columns from left to right in the figure represent gene ID, species name, species family, and expression level, respectively. Interspecific and intraspecific whole-genome duplication events are marked in the upper left corner of the figure. Bootstrap values have been labeled on the phylogenetic tree, which indicate the credibility of the branches. The query genes used in *A. thaliana* have been marked on the tree. The right side of the figure shows the expression data of duplicated genes in dark-red G*. elata*, covering the following developmental stages: TMMM (immature tuber), TMBM (developing tuber), TMJM (mature tuber), TMGS (capsule), and TMHJ (inflorescence). Each stage has three biological replicates, and the data have been averaged.

**Figure S37.** A phylogenetic tree of *ANT* genes from 54 land plant species. The involved plants include 54 plants such as *Gastrodia* and *Dendrobium* (Supplementary Table 20). The vertical columns from left to right in the figure represent gene ID, species name, species family, and expression level, respectively. Interspecific and intraspecific whole-genome duplication events are marked in the upper left corner of the figure. Bootstrap values have been labeled on the phylogenetic tree, which indicate the credibility of the branches. The query genes used in *A. thaliana* have been marked on the tree. The right side of the figure shows the expression data of duplicated genes in dark-red G*. elata*, covering the following developmental stages: TMMM (immature tuber), TMBM (developing tuber), TMJM (mature tuber), TMGS (capsule), and TMHJ (inflorescence). Each stage has three biological replicates, and the data have been averaged.

**Figure S38.** A phylogenetic tree of *PRO* genes from 54 land plant species. The involved plants include 54 plants such as *Gastrodia* and *Dendrobium* (Supplementary Table 20). The vertical columns from left to right in the figure represent gene ID, species name, species family, and expression level, respectively. Interspecific and intraspecific whole-genome duplication events are marked in the upper left corner of the figure. Bootstrap values have been labeled on the phylogenetic tree, which indicate the credibility of the branches. The query genes used in *A. thaliana* have been marked on the tree. The right side of the figure shows the expression data of duplicated genes in dark-red G*. elata*, covering the following developmental stages: TMMM (immature tuber), TMBM (developing tuber), TMJM (mature tuber), TMGS (capsule), and TMHJ (inflorescence). Each stage has three biological replicates, and the data have been averaged.

**Figure S39.** A phylogenetic tree of *AMT* genes from 54 land plant species. The involved plants include 54 plants such as *Gastrodia* and *Dendrobium* (Supplementary Table 20). The vertical columns from left to right in the figure represent gene ID, species name, species family, and expression level, respectively. Interspecific and intraspecific whole-genome duplication events are marked in the upper left corner of the figure. Bootstrap values have been labeled on the phylogenetic tree, which indicate the credibility of the branches. The query genes used in *A. thaliana* have been marked on the tree. The right side of the figure shows the expression data of duplicated genes in dark-red G*. elata*, covering the following developmental stages: TMMM (immature tuber), TMBM (developing tuber), TMJM (mature tuber), TMGS (capsule), and TMHJ (inflorescence). Each stage has three biological replicates, and the data have been averaged.

**Figure S40.** A phylogenetic tree of *Argl* genes from 54 land plant species. The involved plants include 54 plants such as *Gastrodia* and *Dendrobium* (Supplementary Table 20). The vertical columns from left to right in the figure represent gene ID, species name, species family, and expression level, respectively. Interspecific and intraspecific whole-genome duplication events are marked in the upper left corner of the figure. Bootstrap values have been labeled on the phylogenetic tree, which indicate the credibility of the branches. The query genes used in *A. thaliana* have been marked on the tree. The right side of the figure shows the expression data of duplicated genes in dark-red G*. elata*, covering the following developmental stages: TMMM (immature tuber), TMBM (developing tuber), TMJM (mature tuber), TMGS (capsule), and TMHJ (inflorescence). Each stage has three biological replicates, and the data have been averaged.

**Figure S41.** A phylogenetic tree of *Urease* genes from 54 land plant species. The involved plants include 54 plants such as *Gastrodia* and *Dendrobium* (Supplementary Table 20). The vertical columns from left to right in the figure represent gene ID, species name, species family, and expression level, respectively. Interspecific and intraspecific whole-genome duplication events are marked in the upper left corner of the figure. Bootstrap values have been labeled on the phylogenetic tree, which indicate the credibility of the branches. The query genes used in *A. thaliana* have been marked on the tree. The right side of the figure shows the expression data of duplicated genes in dark-red G*. elata*, covering the following developmental stages: TMMM (immature tuber), TMBM (developing tuber), TMJM (mature tuber), TMGS (capsule), and TMHJ (inflorescence). Each stage has three biological replicates, and the data have been averaged.

**Figure S42.** A phylogenetic tree of *NIA* genes from 54 land plant species. The involved plants include 54 plants such as *Gastrodia* and *Dendrobium* (Supplementary Table 20). The vertical columns from left to right in the figure represent gene ID, species name, species family, and expression level, respectively. Interspecific and intraspecific whole-genome duplication events are marked in the upper left corner of the figure. Bootstrap values have been labeled on the phylogenetic tree, which indicate the credibility of the branches. The query genes used in *A. thaliana* have been marked on the tree. The right side of the figure shows the expression data of duplicated genes in dark-red G*. elata*, covering the following developmental stages: TMMM (immature tuber), TMBM (developing tuber), TMJM (mature tuber), TMGS (capsule), and TMHJ (inflorescence). Each stage has three biological replicates, and the data have been averaged.

**Figure S43.** A phylogenetic tree of *NIR* genes from 54 land plant species. The involved plants include 54 plants such as *Gastrodia* and *Dendrobium* (Supplementary Table 20). The vertical columns from left to right in the figure represent gene ID, species name, species family, and expression level, respectively. Interspecific and intraspecific whole-genome duplication events are marked in the upper left corner of the figure. Bootstrap values have been labeled on the phylogenetic tree, which indicate the credibility of the branches. The query genes used in *A. thaliana* have been marked on the tree. The right side of the figure shows the expression data of duplicated genes in dark-red G*. elata*, covering the following developmental stages: TMMM (immature tuber), TMBM (developing tuber), TMJM (mature tuber), TMGS (capsule), and TMHJ (inflorescence). Each stage has three biological replicates, and the data have been averaged.

**Figure S44.** A phylogenetic tree of *NRT* genes from 54 land plant species. The involved plants include 54 plants such as *Gastrodia* and *Dendrobium* (Supplementary Table 20). The vertical columns from left to right in the figure represent gene ID, species name, species family, and expression level, respectively. Interspecific and intraspecific whole-genome duplication events are marked in the upper left corner of the figure. Bootstrap values have been labeled on the phylogenetic tree, which indicate the credibility of the branches. The query genes used in *A. thaliana* have been marked on the tree. The right side of the figure shows the expression data of duplicated genes in dark-red G*. elata*, covering the following developmental stages: TMMM (immature tuber), TMBM (developing tuber), TMJM (mature tuber), TMGS (capsule), and TMHJ (inflorescence). Each stage has three biological replicates, and the data have been averaged.

**Figure S45.** A phylogenetic tree of *OPT* genes from 54 land plant species. The involved plants include 54 plants such as *Gastrodia* and *Dendrobium* (Supplementary Table 20). The vertical columns from left to right in the figure represent gene ID, species name, species family, and expression level, respectively. Interspecific and intraspecific whole-genome duplication events are marked in the upper left corner of the figure. Bootstrap values have been labeled on the phylogenetic tree, which indicate the credibility of the branches. The query genes used in *A. thaliana* have been marked on the tree. The right side of the figure shows the expression data of duplicated genes in dark-red G*. elata*, covering the following developmental stages: TMMM (immature tuber), TMBM (developing tuber), TMJM (mature tuber), TMGS (capsule), and TMHJ (inflorescence). Each stage has three biological replicates, and the data have been averaged.

**Figure S46.** A phylogenetic tree of *GS* genes from 54 land plant species. The involved plants include 54 plants such as *Gastrodia* and *Dendrobium* (Supplementary Table 20). The vertical columns from left to right in the figure represent gene ID, species name, species family, and expression level, respectively. Interspecific and intraspecific whole-genome duplication events are marked in the upper left corner of the figure. Bootstrap values have been labeled on the phylogenetic tree, which indicate the credibility of the branches. The query genes used in *A. thaliana* have been marked on the tree. The right side of the figure shows the expression data of duplicated genes in dark-red G*. elata*, covering the following developmental stages: TMMM (immature tuber), TMBM (developing tuber), TMJM (mature tuber), TMGS (capsule), and TMHJ (inflorescence). Each stage has three biological replicates, and the data have been averaged.

**Figure S47.** A phylogenetic tree of *GLU* genes from 54 land plant species. The involved plants include 54 plants such as *Gastrodia* and *Dendrobium* (Supplementary Table 20). The vertical columns from left to right in the figure represent gene ID, species name, species family, and expression level, respectively. Interspecific and intraspecific whole-genome duplication events are marked in the upper left corner of the figure. Bootstrap values have been labeled on the phylogenetic tree, which indicate the credibility of the branches. The query genes used in *A. thaliana* have been marked on the tree. The right side of the figure shows the expression data of duplicated genes in dark-red G*. elata*, covering the following developmental stages: TMMM (immature tuber), TMBM (developing tuber), TMJM (mature tuber), TMGS (capsule), and TMHJ (inflorescence). Each stage has three biological replicates, and the data have been averaged.

**Figure S48.** A phylogenetic tree of *FAX* genes from 54 land plant species. The involved plants include 54 plants such as *Gastrodia* and *Dendrobium* (Supplementary Table 20). The vertical columns from left to right in the figure represent gene ID, species name, species family, and expression level, respectively. Interspecific and intraspecific whole-genome duplication events are marked in the upper left corner of the figure. Bootstrap values have been labeled on the phylogenetic tree, which indicate the credibility of the branches. The query genes used in *A. thaliana* have been marked on the tree. The right side of the figure shows the expression data of duplicated genes in dark-red G*. elata*, covering the following developmental stages: TMMM (immature tuber), TMBM (developing tuber), TMJM (mature tuber), TMGS (capsule), and TMHJ (inflorescence). Each stage has three biological replicates, and the data have been averaged.

**Figure S49.** A phylogenetic tree of *LACS* genes from 54 land plant species. The involved plants include 54 plants such as *Gastrodia* and *Dendrobium* (Supplementary Table 20). The vertical columns from left to right in the figure represent gene ID, species name, species family, and expression level, respectively. Interspecific and intraspecific whole-genome duplication events are marked in the upper left corner of the figure. Bootstrap values have been labeled on the phylogenetic tree, which indicate the credibility of the branches. The query genes used in *A. thaliana* have been marked on the tree. The right side of the figure shows the expression data of duplicated genes in dark-red G*. elata*, covering the following developmental stages: TMMM (immature tuber), TMBM (developing tuber), TMJM (mature tuber), TMGS (capsule), and TMHJ (inflorescence). Each stage has three biological replicates, and the data have been averaged.

**Figure S50.** A phylogenetic tree of *ABCA* genes from 54 land plant species. The involved plants include 54 plants such as *Gastrodia* and *Dendrobium* (Supplementary Table 20). The vertical columns from left to right in the figure represent gene ID, species name, species family, and expression level, respectively. Interspecific and intraspecific whole-genome duplication events are marked in the upper left corner of the figure. Bootstrap values have been labeled on the phylogenetic tree, which indicate the credibility of the branches. The query genes used in *A. thaliana* have been marked on the tree. The right side of the figure shows the expression data of duplicated genes in dark-red G*. elata*, covering the following developmental stages: TMMM (immature tuber), TMBM (developing tuber), TMJM (mature tuber), TMGS (capsule), and TMHJ (inflorescence). Each stage has three biological replicates, and the data have been averaged.

**Figure S51.** Correlation between gene expression and dN/dS ratio. A: Analysis of the correlation between gene expression levels of copy genes related to carbon and energy transport and dN/dS ratios in dark-red G*. elata*. B: Analysis of the correlation between gene expression levels of copy genes related to nitrogen and energy transport and dN/dS ratios in dark-red G*. elata*. C: Analysis of the correlation between gene expression levels of copy genes related to fatty acid transport and dN/dS ratios in dark-red G*. elata*. For all three panels, the abscissa represents the average expression level of the relevant copy genes across different developmental stages of dark-red G*. elata*, and the ordinate represents the dN/dS ratio of the corresponding copy genes; a dN/dS ratio > 1 indicates that the gene is under positive selection; different gene families are distinguished by different colors.

## Supplementary Table legends

**Supplementary Table 1.** Statistical summary of genome sequencing data of dark-red G*astrodia elata*.

**Supplementary Table 2.** The *Gastrodia elata*’s information on genomic analysis across multiple research articles.

**Supplementary Table 3.** Genome assembly rate of dark-red G*astrodia elata*.

**Supplementary Table 4.** Genomic base composition statistics for dark-red G*astrodia elata*

**Supplementary Table 5.** BUSCO assessment results of the *Gastrodia* genome.

**Supplementary Table 6.** Summary of CEGMA assessment of the dark-red G*astrodia elata* genome.

**Supplementary Table 7.** Read coverage statistics of the dark-red G*astrodia elata* genome.

**Supplementary Table 8.** Single nucleotide polymorphism data of the dark-red G*astrodia elata* genome.

**Supplementary Table 9.** Summary of repeat masking in the dark red *Gastrodia elata* genome. Repetitive regions were identified using TRF (tandem repeats), RepeatMasker (transposable element–related sequences), and ProteinMask (protein-based repeat-associated regions). The total represents non-redundant masked regions.

**Supplementary Table 10.** Summary of gene functional annotations in dark-red G*astrodia elata*.

**Supplementary Table 11.** GO functional enrichment results of non-core genes associated with substance transport.

**Supplementary Table 12.** GO functional enrichment analysis of non-core genes associated with saccharides.

**Supplementary Table 13.** GO functional enrichment analysis of non-core genes associated with nitrogen compounds.

**Supplementary Table 14.** GO and KEGG functional enrichment analysis of non-core genesassociated with light, and GO enrichment analysis on core genes related to light or photosynthesis.

**Supplementary Table 15.** Summary of the transcriptome sequencing data quality across various color forms of *Gastrodia elata* (n=3).

**Supplementary Table 16.** Relative content of volatile components in different germplasms of *Gastrodia elata*, as determined employing GC-MS.

**Supplementary Table 17.** Summary of SNP and INDEL information across 18 chromosomes from dark red dark-red *Gastrodia elata* genome.

**Supplementary Table 18.** Hyde-out-filtered data of 11 *Gastrodia elata*.

**Supplementary Table 19.** Morphology and flowering periods of five *Gastrodia elata* forms (*Flora of China*)

**Supplementary Table 20.** Genomic data sources for 54 land plant species.

**Supplementary Table 21.** The KEGG functional enrichment of gene families related to contraction and expansion in *Gastrodia elata*.

**Supplementary Table 22.** The GO functional enrichment of gene families related to expansion in *Gastrodia elata*.

**Supplementary Table 23.** The GO functional enrichment of gene families related to contraction in *Gastrodia elata*.

**Supplementary Table 24.** Physicochemical Properties of 26 *Armillaria gallica* proteins. **Supplementary Table 25.** Prediction of interactions between 83 *Armillaria gallica* proteins and the GAFP protein.

**Supplementary Table 26.** GO enrichment analysis of 26 *Armillaria gallica* proteins.

**Supplementary Table 27.** KEGG enrichment analysis of 26 *Armillaria gallica* proteins.

**Supplementary Table 28:** Gene expression levels of gastrodin in different developmental stages of dark-red G*. elata*.

**Supplementary Table 29.** The genomic information used in our study to identify HGT-acquired genes.

**Supplementary Table 30.** Horizontally transferred genes identified by HGTphyloDetect, filtered by AI > 45 and outg_pct > 90.

**Supplementary Table 31.** Candidate horizontally transferred acquired genes identified by AvP, filtered by AI ≥ 0 or AHS ≥ 0, with FastTree phylogenetic support.

**Supplementary Table 32.** Information of horizontally transferred acquired genes identified in this study.

**Supplementary Table 33.** The number of read pairs successfully annotated to symbiotic fungi (*Armillaria spp.* and *Mycena spp.*) in the 17 transcriptome datasets of *G. elata*.

**Supplementary Table 34.** Horizontally acquired genes and transcription factor genes, with their KS values at different node depths.

**Supplementary Table 35.** Sequence characteristics of HGT-acquired genes versus transcription factor genes.

**Supplementary Table 36.** Expression profiles (FPKM) of 3 horizontally transferred acquired genes across developmental stages in dark-red G*. elata* and color variant cultivars.

**Supplementary Table 37.** Key information table of query genes involved in carbohydrate and related metabolism.

**Supplementary Table 38.** Supplementary Table 38: Summary of gene copy numbers for energy metabolism–related genes involved in *Gastrodia elata*–fungus interactions across 47 representative plant genomes.

**Supplementary Table 39.** Gene expression levels and temporal clustering analysis of different gene families in different developmental stages of dark-red G*. elata*.

**Supplementary Table 40.** Quantification of fatty acid contents in *G. elata* at various developmental stages using HPLC (μg/g).

## References

1. Li, J., Zhang, M., Yang, Z., and Li, C. *Botrytis cinerea* causes flower gray mold in *Gastrodia elata* in China. Crop Prot. 2022; 155: 105923.

2. Zhang, G.-Q., Xu, Q., Bian, C., Tsai, W.-C., Yeh, C.-M., Liu, K.-W., Yoshida, K., Zhang, L.-S., Chang, S.-B., and Chen, F. The *Dendrobium catenatum* Lindl. genome sequence provides insights into polysaccharide synthase, floral development and adaptive evolution. Sci. Rep. 2016; 6: 19029.

3. Sun, X., Jia, B., Sun, J., Lin, J., Lu, B., Duan, J., Li, C., Wang, Q., Zhang, X., and Tan, M. *Gastrodia elata* Blume: A review of its mechanisms and functions on cardiovascular systems. Fitoterapia 2023: 105511.

4. Chen, S., Wang, X., Wang, Y., Zhang, G., Song, W., Dong, X., Arnold, M.L., Wang, W., Miao, J., and Chen, W. Improved *de novo* assembly of the achlorophyllous orchid *Gastrodia elata*. Front. Genet. 2020; 11: 580568.

5. Xu, Y., Lei, Y., Su, Z., Zhao, M., Zhang, J., Shen, G., Wang, L., Li, J., Qi, J., and Wu, J. A chromosome-scale *Gastrodia elata* genome and large-scale comparative genomic analysis indicate convergent evolution by gene loss in mycoheterotrophic and parasitic plants. Plant J. 2021; 108: 1609–1623.

6. Bae, E.-K., An, C., Kang, M.-J., Lee, S.-A., Lee, S.J., Kim, K.-T., and Park, E.-J. Chromosome-level genome assembly of the fully mycoheterotrophic orchid Gastrodia elata. G3-Genes, Genomes, Genetics. 2022; 12: jkab433.

7. Yang, J., Li, P., Li, Y., and Xiao, Q. GelFAP v2. 0: an improved platform for Gene functional analysis in *Gastrodia elata*. BMC Genom. 2023; 24: 1–12.

8. Suetsugu, K. Living in the shadows: *Gastrodia* orchids lack functional leaves and open flowers. Plants people planet. 2022; 4: 418–422.

9. Tian, Y., Dong, Q., Ji, Z., Chi, F., Cong, P., and Zhou, Z. Genome-wide identification and analysis of the MADS-box gene family in apple. Gene. 2015; 555: 277–290.

10. Cox, K.D., Layne, D.R., Scorza, R., and Schnabel, G. *Gastrodia* anti-fungal protein from the orchid *Gastrodia elata* confers disease resistance to root pathogens in transgenic tobacco. Planta. 2006; 224 (*Planta*): 1373–1383. 10.1007/s00425-006-0322-0.

11. Jiang, Y., Hu, X., Yuan, Y., Guo, X., Chase, M.W., Ge, S., Li, J., Fu, J., Li, K., and Hao, M.J.B.P.B. The *Gastrodia menghaiensis* (Orchidaceae) genome provides new insights of orchid mycorrhizal interactions. BMC Plant Biol. 2022; 22: 179.

12. Xu, Q., Liu, Y., Wang, X., Gu, H., Chen, Z.J.P.p., and Biochemistry. Purification and characterization of a novel anti-fungal protein from *Gastrodia elata*. Plant Physiol. Biochem. 1998; 36: 899–905.

13. Chen, R., Huangfu, L., Lu, Y., Fang, H., Xu, Y., Li, P., Zhou, Y., Xu, C., Huang, J., and Yang, Z. Adaptive innovation of green plants by horizontal gene transfer. Biotechnol. Adv. 2021; 46: 107671.

14. Graham, L.A., Lougheed, S.C., Ewart, K.V., and Davies, P.L. Lateral transfer of a lectin-like antifreeze protein gene in fishes. PloS one. 2008; 3: e2616.

15. Wang, H., Sun, S., Ge, W., Zhao, L., Hou, B., Wang, K., Lyu, Z., Chen, L., Xu, S., and Guo, J. Horizontal gene transfer of *Fhb7* from fungus underlies *Fusarium* head blight resistance in wheat. Science. 2020; 368: eaba5435.

16. Ludewig, F., and Flügge, U.-I. Role of metabolite transporters in source-sink carbon allocation. Front. Plant Sci. 2013; 4: 231.

17. Camprubi, A., Solari, J., Bonini, P., Garcia-Figueres, F., Colosimo, F., Cirino, V., Lucini, L., and Calvet, C. Plant Performance and Metabolomic Profile of Loquat in Response to Mycorrhizal Inoculation, Armillaria mellea and Their Interaction. Agronomy. 2020; 10: 899.

18. Chen, L., Wang, Y.-C., Qin, L.-Y., He, H.-Y., Yu, X.-L., Yang, M.-Z., and Zhang, H.-B. Dynamics of fungal communities during *Gastrodia elata* growth. BMC Microbiol. 2019; 19: 1–11.

19. Merckx, V. Mycoheterotrophy: the biology of plants living on fungi. Springer Science & Business Media. 2012.

20. Ho, L.H., Lee, Y.I., Hsieh, S.Y., Lin, I.S., Wu, Y.C., Ko, H.Y., Klemens, P.A., Neuhaus, H.E., Chen, Y.M., and Huang, T.P. *Ge*SUT4 mediates sucrose import at the symbiotic interface for carbon allocation of heterotrophic *Gastrodia elata* (Orchidaceae). Plant Cell Environ. 2021; 44: 20–33.

21. Ainsworth, E.A., and Bush, D.R. Carbohydrate export from the leaf: a highly regulated process and target to enhance photosynthesis and productivity. Plant Physiol. 2011; 155: 64–69.

22. Ludewig, F., and Flügge, U.-I.J.F.i.P.S. Role of metabolite transporters in source-sink carbon allocation. Front. Plant Sci. 2013; 4: 231.

23. O’Leary, B., Park, J., and Plaxton, W.C.J.B.J. The remarkable diversity of plant PEPC (phosphoenolpyruvate carboxylase): recent insights into the physiological functions and post-translational controls of non-photosynthetic PEPCs. Biochem. J. 2011; 436: 15–34.

24. Motomura, H., Yukawa, T., Ueno, O., and Kagawa, A. The occurrence of crassulacean acid metabolism in *Cymbidium* (Orchidaceae) and its ecological and evolutionary implications. J. Plant Res. 2008; 121: 163–177.

25. Dong, N.Q., and Lin, H.X. Contribution of phenylpropanoid metabolism to plant development and plant-environment interactions. J. Integr. Plant Biol. 2021; 63: 180–209.

26. Zhang, G.-Q., Liu, K.-W., Li, Z., Lohaus, R., Hsiao, Y.-Y., Niu, S.-C., Wang, J.-Y., Lin, Y.-C., Xu, Q., and Chen, L.-J. The *Apostasia* genome and the evolution of orchids. Nature. 2017; 549: 379–383.

27. Wang, Y., Liang, C., Wu, S., Zhang, X., Tang, J., Jian, G., Jiao, G., Li, F., and Chu, C. Significant Improvement of Cotton *Verticillium* Wilt Resistance by Manipulating the Expression of *Gastrodia* Antifungal Proteins. Mol. Plant. 2016; 9: 1436–1439. 10.1016/j.molp.2016.06.013.

28. Wang, Y., Liang, C., Wu, S., Jian, G., Zhang, X., Zhang, H., Tang, J., Li, J., Jiao, G., Li, F., and Chu, C. Vascular-specific expression of *Gastrodia* antifungal protein gene significantly enhanced cotton *Verticillium* wilt resistance. Plant Biotechnol. J. 2020; 18: 1498–1500. 10.1111/pbi.13308.

29. Salehi, N., and Peng, C.-A. Purification of CD47-streptavidin fusion protein from bacterial lysate using biotin-agarose affinity chromatography. Biotechnol. Prog. 2016; 32: 949–958.

30. Mohammed, H., Taylor, C., Brown, G.D., Papachristou, E.K., Carroll, J.S., and D’santos, C.S. Rapid immunoprecipitation mass spectrometry of endogenous proteins (RIME) for analysis of chromatin complexes. Nat. Protoc. 2016; 11: 316–326.

31. Turriziani, B., Garcia-Munoz, A., Pilkington, R., Raso, C., Kolch, W., and Von Kriegsheim, A. On-beads digestion in conjunction with data-dependent mass spectrometry:a shortcut to quantitative and dynamic interaction proteomics. Biol. 2014; 3: 320–332.

32. Gasteiger, E., Hoogland, C., Gattiker, A., Duvaud, S.e., Wilkins, M.R., Appel, R.D., and Bairoch, A. Protein Identification and Analysis Tools on the ExPASy Server. The Proteomics Protocols Handbook. 2005: 571–607. 10.1385/1-59259-890-0:571.

33. Yugandhar, K., and Gromiha, M.M. Protein-protein binding affinity prediction from amino acid sequence. Bioinformatics. 2014; 30: 3583–3589. 10.1093/bioinformatics/btu580.

34. Nikam, R., Yugandhar, K., and Michael Gromiha, M. Discrimination and Prediction of Protein-Protein Binding Affinity Using Deep Learning Approach. Intelligent Computing Theories and Application. 2018: 809–815.

35. Commission, C.P. Pharmacopoeia of the People’s Republic of China 2020. China Medical Science Press. 2020; 59 (59).

36. Yin, H., Hu, T., Zhuang, Y., and Liu, T. Metabolic engineering of *Saccharomyces cerevisiae* for high-level production of gastrodin from glucose. Microb. Cell Factories. 2020; 19: 1–12.

37. Tsai, C.-C., Wu, K.-M., Chiang, T.-Y., Huang, C.-Y., Chou, C.-H., Li, S.-J., and Chiang, Y.-C. Comparative transcriptome analysis of *Gastrodia elata* (Orchidaceae) in response to fungus symbiosis to identify gastrodin biosynthesis-related genes. BMC Genom. 2016; 17: 1–16.

38. Becker, A., and Theißen, G. The major clades of MADS-box genes and their role in the development and evolution of flowering plants. Mol. Phylogenet. 2003; 29: 464–489.

39. Parenicova, L., de Folter, S., Kieffer, M., Horner, D.S., Favalli, C., Busscher, J., Cook, H.E., Ingram, R.M., Kater, M.M., and Davies, B. Molecular and phylogenetic analyses of the complete MADS-box transcription factor family in Arabidopsis: new openings to the MADS world. Plant Cell. 2003; 15: 1538–1551.

40. Theißen, G., Melzer, R., and Rümpler, F. MADS-domain transcription factors and the floral quartet model of flower development: linking plant development and evolution. Development. 2016; 143: 3259–3271.

41. Adamczyk B J, L.S.M.D., Fernandez D E. The MADS domain factors AGL15 and AGL18 act redundantly as repressors of the floral transition in Arabidopsis. Plant J. 2007; 50: 1007–1019.

42. Tapia-López R, G.-P.B., Dubrovsky J G. An AGAMOUS-related MADS-box gene, XAL1 (AGL12), regulates root meristem cell proliferation and flowering transition in Arabidopsis. Plant Physiol. 2008; 146: 1182–1192.

43. Dreni L, Z.D. Flower development: the evolutionary history and functions of the *AGL6* subfamily MADS-box genes. J. Exp. Bot. 2016; 67: 1625–1638.

44. Liu, J.-J., Yang, X.-Q., Li, Z.-Y., Miao, J.-Y., Li, S.-B., Zhang, W.-P., Lin, Y.-C., and Lin, L.-B. The role of symbiotic fungi in the life cycle of *Gastrodia elata* Blume (Orchidaceae): A comprehensive review. Front. Plant Sci. 2024; 14: 1309038.

45. Suetsugu, K., Matsubayashi, J., and Tayasu, I. Some mycoheterotrophic orchids depend on carbon from dead wood: novel evidence from a radiocarbon approach. New Phytol. 2020; 227: 1519–1529.

46. Smith, S.E. Carbohydrate translocation in orchid mycorrhizas. New Phytol. 1967; 66: 371–378.

47. Ponert, J., Šoch, J., Vosolsobě, S., Čiháková, K., and Lipavská, H. Integrative study supports the role of trehalose in carbon transfer from fungi to mycotrophic orchid. Front. Plant Sci. 2021; 12: 793876.

48. Büttner, M. The *Arabidopsis* sugar transporter (AtSTP) family: an update. Plant Biol. 2010; 12: 35–41.

49. Büttner, M., Truernit, E., Baier, K., Scholz-Starke, J., Sontheim, M., Lauterbach, C., Huss, V., and Sauer, N. AtSTP3, a green leaf-specific, low affinity monosaccharide-H^+^ symporter of *Arabidopsis thaliana*. Plant Cell Environ. 2000; 23: 175–184.

50. Poschet, G., Hannich, B., and Büttner, M. Identification and characterization of AtSTP14, a novel galactose transporter from Arabidopsis. Plant Cell Physiol. 2010; 51: 1571–1580.

51. Büttner, M. The monosaccharide transporter (-like) gene family in *Arabidopsis*. FEBS Lett. 2007; 581: 2318–2324.

52. Amor, Y., Haigler, C.H., Johnson, S., Wainscott, M., and Delmer, D.P. A membrane-associated form of sucrose synthase and its potential role in synthesis of cellulose and callose in plants. Proc. Natl. Acad. Sci. U.S.A. 1995; 92: 9353–9357.

53. Yao, D., Gonzales-Vigil, E., and Mansfield, S.D. Arabidopsis sucrose synthase localization indicates a primary role in sucrose translocation in phloem. J. Exp. Bot. 2020; 71: 1858–1869.

54. Yao, X., Nie, J., Bai, R., and Sui, X. Amino acid transporters in plants: Identification and function. Plants. 2020; 9: 972.

55. Rentsch, D., Schmidt, S., and Tegeder, M. Transporters for uptake and allocation of organic nitrogen compounds in plants. FEBS Lett. 2007; 581: 2281–2289.

56. Wang, R., Guan, P., Chen, M., Xing, X., Zhang, Y., and Crawford, N.M. Multiple regulatory elements in the Arabidopsis *NIA1* promoter act synergistically to form a nitrate enhancer. Plant Physiol. 2010; 154: 423–432.

57. Costa-Broseta, Á., Castillo, M., and León, J. Nitrite reductase 1 is a target of nitric oxide-mediated post-translational modifications and controls nitrogen flux and growth in Arabidopsis. Int. J. Mol. Sci. 2020; 21: 7270.

58. Cruz, C., Egsgaard, H., Trujillo, C., Ambus, P., Requena, N., Martins-Loução, M.A., and Jakobsen, I. Enzymatic evidence for the key role of arginine in nitrogen translocation by arbuscular mycorrhizal fungi. Plant Physiol. 2007; 144: 782–792.

59. Gazzarrini, S., Lejay, L., Gojon, A., Ninnemann, O., Frommer, W.B., and von Wirén, N. Three functional transporters for constitutive, diurnally regulated, and starvation-induced uptake of ammonium into Arabidopsis roots. Plant Cell. 1999; 11: 937–947.

60. Bu, Y., Kou, J., Sun, B., Takano, T., and Liu, S. Adverse effect of urease on salt stress during seed germination in *Arabidopsis thaliana*. FEBS Lett. 2015; 589: 1308–1313.

61. Yuan, Y., Jin, X., Liu, J., Zhao, X., Zhou, J., Wang, X., Wang, D., Lai, C., Xu, W., and Huang, J. The *Gastrodia elata* genome provides insights into plant adaptation to heterotrophy. Nat. Commun. 2018; 9: 1615.

62. Koh, S., Wiles, A.M., Sharp, J.S., Naider, F.R., Becker, J.M., and Stacey, G. An oligopeptide transporter gene family in Arabidopsis. Plant Physiol*. .* 2002; 128: 21–29.

63. Nam, J.-W., Lee, H.G., Do, H., Kim, H.U., and Seo, P.J. Transcriptional regulation of triacylglycerol accumulation in plants under environmental stress conditions. J. Exp. Bot. 2022; 73: 2905–2917.

64. Li, N., Gügel, I.L., Giavalisco, P., Zeisler, V., Schreiber, L., Soll, J., and Philippar, K. FAX1, a novel membrane protein mediating plastid fatty acid export. PLoS Biol. 2015; 13: e1002053.

65. Kim, S., Yamaoka, Y., Ono, H., Kim, H., Shim, D., Maeshima, M., Martinoia, E., Cahoon, E.B., Nishida, I., and Lee, Y. AtABCA9 transporter supplies fatty acids for lipid synthesis to the endoplasmic reticulum. Proc. Natl Acad. Sci. USA. 2013; 110: 773–778.

66. Ayaz, A., Saqib, S., Huang, H., Zaman, W., Lü, S., and Zhao, H. Genome-wide comparative analysis of long-chain acyl-CoA synthetases (LACSs) gene family: A focus on identification, evolution and expression profiling related to lipid synthesis. Plant Physiol. Biochem. 2021; 161: 1–11.

67. Jiang, Y., Wang, W., Xie, Q., Liu, N., Liu, L., Wang, D., Zhang, X., Yang, C., Chen, X., and Tang, D. Plants transfer lipids to sustain colonization by mutualistic mycorrhizal and parasitic fungi. Science 2017; 356: 1172–1175.

68. Hu Z, H.Q. Induction and accumulation of the antifungal protein in Gastrodia elata. Plant Divers. 1994; 16: 1.

69. Zhang, X., Zhang, S., Zhao, Q., Ming, R., and Tang, H. Assembly of allele-aware, chromosomal-scale autopolyploid genomes based on Hi-C data. Nat. Plants. 2019; 5: 833–845.

70. Robinson, J.T., Turner, D., Durand, N.C., Thorvaldsdóttir, H., Mesirov, J.P., and Aiden, E.L. Juicebox. js provides a cloud-based visualization system for Hi-C data. Cell Syst. 2018; 6: 256–258. e251.

71. Manni, M., Berkeley, M.R., Seppey, M., Simão, F.A., and Zdobnov, E.M. BUSCO update: novel and streamlined workflows along with broader and deeper phylogenetic coverage for scoring of eukaryotic, prokaryotic, and viral genomes. Mol. Biol. Evol. 2021; 38: 4647–4654.

72. Parra, G., Bradnam, K., and Korf, I. CEGMA: a pipeline to accurately annotate core genes in eukaryotic genomes. Bioinformatics. 2007; 23: 1061–1067.

73. Li, H., and Durbin, R. Fast and accurate short read alignment with Burrows–Wheeler transform. Bioinformatics. 2009; 25: 1754–1760.

74. Rhie, A., Walenz, B.P., Koren, S., and Phillippy, A.M. Merqury: reference-free quality, completeness, and phasing assessment for genome assemblies. Genome Biol. 2020; 21: 245. 10.1186/s13059-020-02134-9.

75. Jurka, J., Kapitonov, V.V., Pavlicek, A., Klonowski, P., Kohany, O., and Walichiewicz, J. Repbase Update, a database of eukaryotic repetitive elements. Cytogenet. Genome Res. 2005; 110: 462–467.

76. Altschul, S.F., Madden, T.L., Schäffer, A.A., Zhang, J., Zhang, Z., Miller, W., and Lipman, D.J. Gapped BLAST and PSI-BLAST: a new generation of protein database search programs. Nucleic Acids Res. 1997; 25: 3389–3402.

77. Birney, E., Clamp, M., and Durbin, R. GeneWise and genomewise. Genome Res. 2004; 14: 988–995.

78. Nachtweide, S., and Stanke, M. Multi-genome annotation with AUGUSTUS. Methods Mol Biol. 2019; 1962: 139–160.

79. Blanco, E., Parra, G., and Guigó, R. Using Geneid to identify genes. Hoboken. Curr Protoc Bioinformatics. 2007: Chapter 4:Unit 4.3.

80. Burge, C., and Karlin, S. Prediction of complete gene structures in human genomic DNA. J Mol Biol. 1997; 268: 78–94.

81. Majoros, W.H., Pertea, M., and Salzberg, S.L. TigrScan and GlimmerHMM: two open source ab initio eukaryotic gene-finders. Bioinformatics. 2004; 20: 2878–2879.

82. Korf, I. Gene finding in novel genomes. BMC Bioinformatics. 2004; 5: 1–9.

83. Kim, D., Paggi, J.M., Park, C., Bennett, C., and Salzberg, S.L. Graph-based genome alignment and genotyping with HISAT2 and HISAT-genotype. Nat Biotechnol*. .* 2019; 37: 907–915.

84. Trapnell, C., Pachter, L., and Salzberg, S.L. TopHat: discovering splice junctions with RNA-Seq. Bioinformatics. 2009; 25: 1105–1111.

85. Pertea, M., Pertea, G.M., Antonescu, C.M., Chang, T.-C., Mendell, J.T., and Salzberg, S.L. StringTie enables improved reconstruction of a transcriptome from RNA-seq reads. Nat. Biotechnol. 2015; 33: 290–295.

86. Ghosh, S., and Chan, C.-K.K. Analysis of RNA-Seq data using TopHat and Cufflinks. Methods Mol. Biol. 2016; 1374: 339–361.

87. Jones, P., Binns, D., Chang, H.-Y., Fraser, M., Li, W., McAnulla, C., McWilliam, H., Maslen, J., Mitchell, A., and Nuka, G. InterProScan 5: genome-scale protein function classification. Bioinformatics. 2014; 30: 1236–1240.

88. Lowe, T.M., and Eddy, S.R. tRNAscan-SE: a program for improved detection of transfer RNA genes in genomic sequence. Nucleic Acids Res. 1997; 25: 955–964.

89. Nawrocki, E.P., and Eddy, S.R. Infernal 1.1: 100-fold faster RNA homology searches. Bioinformatics. 2013; 29: 2933–2935.

90. Smit, A.F., Hubley, R., and Green, P. RepeatModeler Open-1.0. Available online: https://www.repeatmasker.org/RepeatModeler. 2008.

91. Cantarel, B.L., Korf, I., Robb, S.M., Parra, G., Ross, E., Moore, B., Holt, C., Alvarado, A.S., and Yandell, M. MAKER: an easy-to-use annotation pipeline designed for emerging model organism genomes. Genome Res. 2008; 18: 188–196.

92. Cai, J., Liu, X., Vanneste, K., Proost, S., Tsai, W.-C., Liu, K.-W., Chen, L.-J., He, Y., Xu, Q., and Bian, C. The genome sequence of the orchid *Phalaenopsis equestris*. Nat. Genet. 2015; 47: 65–72.

93. Keilwagen, J., Hartung, F., and Grau, J. GeMoMa: homology-based gene prediction utilizing intron position conservation and RNA-seq data. Gene prediction: Methods and protocols. 2019: 161–177.

94. Bolger, A.M., Lohse, M., and Usadel, B. Trimmomatic: a flexible trimmer for Illumina sequence data. Bioinformatics. 2014; 30: 2114–2120.

95. Luo, R., Liu, B., Xie, Y., Li, Z., Huang, W., Yuan, J., He, G., Chen, Y., Pan, Q., and Liu, Y. SOAPdenovo2: an empirically improved memory-efficient short-read de novo assembler. GigaScience. 2012; 1: 2047-2217X-2041-2018.

96. Grabherr, M.G., Haas, B.J., Yassour, M., Levin, J.Z., Thompson, D.A., Amit, I., Adiconis, X., Fan, L., Raychowdhury, R., and Zeng, Q. Full-length transcriptome assembly from RNA-Seq data without a reference genome. Nat. Biotechnol. 2011; 29: 644–652.

97. Haas, B.J., Papanicolaou, A., Yassour, M., Grabherr, M., Blood, P.D., Bowden, J., Couger, M.B., Eccles, D., Li, B., and Lieber, M. *De novo* transcript sequence reconstruction from RNA-seq using the Trinity platform for reference generation and analysis. Nat. Protoc. 2013; 8: 1494–1512.

98. Fu, L., Niu, B., Zhu, Z., Wu, S., and Li, W.J.B. CD-HIT: accelerated for clustering the next-generation sequencing data. Bioinformatics. 2012; 28: 3150–3152.

99. Camacho, C., Coulouris, G., Avagyan, V., Ma, N., Papadopoulos, J., Bealer, K., and Madden, T.L. BLAST+: architecture and applications. BMC Bioinform. 2009; 10: 1–9.

100. Simão, F.A., Waterhouse, R.M., Ioannidis, P., Kriventseva, E.V., and Zdobnov, E.M.J.B. BUSCO: assessing genome assembly and annotation completeness with single-copy orthologs. Bioinformatics. 2015; 31: 3210–3212.

101. Emms, D.M., and Kelly, S. OrthoFinder: phylogenetic orthology inference for comparative genomics. Genome Biol. 2019; 20: 1–14.

102. McKenna, A., Hanna, M., Banks, E., Sivachenko, A., Cibulskis, K., Kernytsky, A., Garimella, K., Altshuler, D., Gabriel, S., and Daly, M. The Genome Analysis Toolkit: a MapReduce framework for analyzing next-generation DNA sequencing data. Genome Res. 2010; 20: 1297–1303.

103. Purcell, S., Neale, B., Todd-Brown, K., Thomas, L., Ferreira, M.A., Bender, D., Maller, J., Sklar, P., De Bakker, P.I., and Daly, M.J. PLINK: a tool set for whole-genome association and population-based linkage analyses. Am. J. Hum. Genet. 2007; 81: 559–575.

104. Ortiz, E.M. vcf2phylip v2.0: convert a VCF matrix into several matrix formats for phylogenetic analysis. Available from: https://github.com/edgardomortiz/vcf2phylip. 2019.

105. Nguyen, L.-T., Schmidt, H.A., Von Haeseler, A., and Minh, B.Q. IQ-TREE: a fast and effective stochastic algorithm for estimating maximum-likelihood phylogenies. Mol. Biol. Evol. 2015; 32: 268–274.

106. Kalyaanamoorthy, S., Minh, B.Q., Wong, T.K., Von Haeseler, A., and Jermiin, L.S. ModelFinder: fast model selection for accurate phylogenetic estimates. Nat. Methods. 2017; 14: 587–589.

107. Blischak, P.D., Chifman, J., Wolfe, A.D., and Kubatko, L.S. HyDe: a Python package for genome-scale hybridization detection. Syst. Biol. 2018; 67: 821–829.

108. Wang, Y., Tang, H., DeBarry, J.D., Tan, X., Li, J., Wang, X., Lee, T.-h., Jin, H., Marler, B., and Guo, H. *MCScanX*: a toolkit for detection and evolutionary analysis of gene synteny and collinearity. Nucleic Acids Res. 2012; 40: e49–e49.

109. Wang, X., Shi, X., Li, Z., Zhu, Q., Kong, L., Tang, W., Ge, S., and Luo, J. Statistical inference of chromosomal homology based on gene colinearity and applications to *Arabidopsis* and rice. BMC Bioinform. 2006; 7: 1–13.

110. Sun, P., Jiao, B., Yang, Y., Shan, L., Li, T., Li, X., Xi, Z., Wang, X., and Liu, J. WGDI: A user-friendly toolkit for evolutionary analyses of whole-genome duplications and ancestral karyotypes. Mol. Plant. 2022; 15: 1841–1851.

111. Yang, Z. PAML 4: phylogenetic analysis by maximum likelihood. Mol. Biol. Evol. 2007; 24: 1586–1591.

112. Zhao, Y., Zhang, R., Jiang, K.-W., Qi, J., Hu, Y., Guo, J., Zhu, R., Zhang, T., Egan, A.N., and Yi, T.-S. Nuclear phylotranscriptomics and phylogenomics support numerous polyploidization events and hypotheses for the evolution of rhizobial nitrogen-fixing symbiosis in Fabaceae. Mol. Plant. 2021; 14: 748–773.

113. Van Dongen, S. Graph clustering via a discrete uncoupling process. SIAM J. Matrix Anal. Appl. 2008; 30: 121–141.

114. Katoh, K., and Standley, D.M. MAFFT multiple sequence alignment software version 7: improvements in performance and usability. Mol. Biol. Evol. 2013; 30: 772–780.

115. Suyama, M., Torrents, D., and Bork, P. PAL2NAL: robust conversion of protein sequence alignments into the corresponding codon alignments. Nucleic Acids Res. 2006; 34: W609–W612.

116. Capella-Gutiérrez, S., Silla-Martínez, J.M., and Gabaldón, T. trimAl: a tool for automated alignment trimming in large-scale phylogenetic analyses. Bioinformatics. 2009; 25: 1972–1973.

117. Iles, W.J., Smith, S.Y., Gandolfo, M.A., and Graham, S.W. Monocot fossils suitable for molecular dating analyses. Bot. J. Linn. Soc. 2015; 178: 346–374.

118. Wu, Y., You, H.-L., and Li, X.-Q. Dinosaur-associated Poaceae epidermis and phytoliths from the Early Cretaceous of China. Natl. Sci. Rev. 2018; 5: 721–727.

119. Magallón, S., Gómez-Acevedo, S., Sánchez-Reyes, L.L., and Hernández-Hernández, T. A metacalibrated time-tree documents the early rise of flowering plant phylogenetic diversity. New Phytol. 2015; 207: 437–453.

120. Morris, J.L., Puttick, M.N., Clark, J.W., Edwards, D., Kenrick, P., Pressel, S., Wellman, C.H., Yang, Z., Schneider, H., and Donoghue, P.C. The timescale of early land plant evolution. Proc. Natl Acad. Sci. USA. 2018; 115: E2274–E2283.

121. Barba-Montoya, J., Dos Reis, M., Schneider, H., Donoghue, P.C., and Yang, Z. Constraining uncertainty in the timescale of angiosperm evolution and the veracity of a Cretaceous Terrestrial Revolution. New Phytol. 2018; 218: 819–834.

122. Buchfink, B., Xie, C., and Huson, D.H. Fast and sensitive protein alignment using DIAMOND. Nat. Methods. 2015; 12: 59–60.

123. Zhou, S., Chen, Y., Guo, C., and Qi, J. PhyloMCL: accurate clustering of hierarchical orthogroups guided by phylogenetic relationship and inference of polyploidy events. Methods Ecol. Evol. 2020; 11: 943–954.

124. Chernomor, O., von Haeseler, A., and Minh, B.Q. Terrace Aware Data Structure for Phylogenomic Inference from Supermatrices. Syst. Biol. 2016; 65: 997–1008. 10.1093/sysbio/syw037.

125. Huerta-Cepas, J., Serra, F., and Bork, P. ETE 3: reconstruction, analysis, and visualization of phylogenomic data. Mol. Biol. Evol. 2016; 33: 1635–1638.

126. Wingett, S.W., and Andrews, S. FastQ Screen: A tool for multi-genome mapping and quality control. F1000research. 2018; 7.

127. Kim, D., Langmead, B., and Salzberg, S.L.J.N.m. HISAT: a fast spliced aligner with low memory requirements. Nat. Methods. 2015; 12: 357–360.

128. Liao, Y., Smyth, G.K., and Shi, W. The R package Rsubread is easier, faster, cheaper and better for alignment and quantification of RNA sequencing reads. Nucleic Acids Res. 2019; 47: e47–e47.

129. Love, M.I., Huber, W., and Anders, S. Moderated estimation of fold change and dispersion for RNA-seq data with DESeq2. Genome Biol. 2014; 15: 1–21.

130. Letunic, I., and Bork, P. Interactive Tree Of Life (iTOL) v5: an online tool for phylogenetic tree display and annotation. Nucleic Acids Res. 2021; 49: W293–W296.

131. Tyanova, S., Temu, T., and Cox, J. The MaxQuant computational platform for mass spectrometry-based shotgun proteomics. Nat. Protoc. 2016; 11: 2301–2319.

132. Madeira, F., Madhusoodanan, N., Lee, J., Eusebi, A., Niewielska, A., Tivey, A.R., Lopez, R., and Butcher, S. The EMBL-EBI Job Dispatcher sequence analysis tools framework in 2024. Nucleic Acids Res. 2024: gkae241.

133. Bateman, A., Coin, L., Durbin, R., Finn, R.D., Hollich, V., Griffiths-Jones, S., Khanna, A., Marshall, M., Moxon, S., and Sonnhammer, E.L. The Pfam protein families database. Nucleic Acids Res. 2004; 32: D138–D141.

134. Yuan, L., Lu, H., Li, F., Nielsen, J., and Kerkhoven, E.J. HGTphyloDetect: facilitating the identification and phylogenetic analysis of horizontal gene transfer. Brief. Bioinform. 2023; 24: bbad035.

135. Koutsovoulos, G.D., Granjeon Noriot, S., Bailly-Bechet, M., Danchin, E.G., and Rancurel, C. AvP: a software package for automatic phylogenetic detection of candidate horizontal gene transfers. PLoS Comput. Biol. 2022; 18: e1010686.

136. One Thousand Plant Transcriptomes Initiative. One thousand plant transcriptomes and the phylogenomics of green plants. Nature. 2019; 574: 679–685.

137. Price, M.N., Dehal, P.S., and Arkin, A.P. FastTree 2 – Approximately Maximum-Likelihood Trees for Large Alignments. PLoS One. 2010; 5: e9490. 10.1371/journal.pone.0009490.

138. Bushnell, B. BBMap: a fast, accurate, splice-aware aligner. 2014.

139. Shannon, P., Markiel, A., Ozier, O., Baliga, N.S., Wang, J.T., Ramage, D., Amin, N., Schwikowski, B., and Ideker, T. Cytoscape: a software environment for integrated models of biomolecular interaction networks. Genome Res. 2003; 13: 2498–2504.

140. Bindea, G., Mlecnik, B., Hackl, H., Charoentong, P., Tosolini, M., Kirilovsky, A., Fridman, W.-H., Pagès, F., Trajanoski, Z., and Galon, J. ClueGO: a Cytoscape plug-in to decipher functionally grouped gene ontology and pathway annotation networks. Bioinformatics. 2009; 25: 1091–1093.

141. Reinders, A., Sivitz, A.B., and Ward, J.M. Evolution of plant sucrose uptake transporters. Front. plant sci. 2012; 3: 20960.

142. Chen, D., Zhang, T., Chen, Y., Ma, H., and Qi, J. Tree2GD: a phylogenomic method to detect large-scale gene duplication events. Bioinformatics. 2022; 38: 5317–5321.

143. Ernst, J., and Bar-Joseph, Z. STEM: a tool for the analysis of short time series gene expression data. BMC bioinformatics. 2006; 7: 1–11.

